# Feedforward and disinhibitory circuits differentially control activity of cortical somatostatin-positive interneurons during behavioral state transitions

**DOI:** 10.1101/2024.01.10.574973

**Authors:** Marcel de Brito Van Velze, Dhanasak Dhanasobhon, Marie Martinez, Annunziato Morabito, Emmanuelle Berthaux, Cibele Martins Pinho, Yann Zerlaut, Nelson Rebola

## Abstract

Local inhibitory neurons (INs), specifically those involved in disinhibitory circuits like somatostatin (SST) and vasoactive intestinal peptide (VIP)-positive INs, are essential for regulating cortical function during behavior. However, it is unclear the mechanisms by which these INs are recruited during active states and whether their activity is consistent across different sensory cortices. We now reveal that in mice, locomotor activity strongly recruits SST-INs in primary somatosensory (S1) but not visual (V1) cortex. This diverse engagement of SST-INs could not be explained by differences in VIP-IN function but was absent in the presence of visual sensory drive suggesting involvement of feedforward sensory pathways. In agreement, inactivating the somatosensory thalamus, but not decreasing VIP-INs activity, significantly reduces the recruitment of SST-INs in S1 by locomotion. Finally, model simulations suggest differences in SST-INs activity can be explained by varying ratios of VIP-driven inhibition and thalamus-driven cortical excitation. Our work suggests that by integrating feedforward activity with neuromodulation, SST-INs play a central role in adapting sensory processing to behavioral states.

---

In the mammalian brain, the cerebral cortex is responsible for complex cognitive functions, including forming a coherent representation of the external world that is crucial for an effective interaction with a constantly changing environment. Importantly, sensation can occur under drastically different behavioral conditions, which have been reported to influence perceptual outcomes^1,2^.

Local neocortical interneurons, in particular those involved in disinhibitory circuits, like vasoactive intestinal peptide (VIP) and somatostatin (SST) expressing interneurons (INs), have been shown to be sensitive to behavior state transitions and to be key contributors to the state-dependent regulation of cortical function^3–9^. In the primary visual cortex (V1), recruitment of VIP-INs and consequent decrease in SST-IN mediated inhibition has been suggested to underlie the gain modulation of visual-evoked responses observed during locomotion^4^. Interestingly, behavioural state-dependent gain modulation measured at the level of pyramidal neurons (PNs) seems to be diverse across sensory cortices ^8,10–16^. In V1, locomotion enhances the responses of PNs to drifting grating stimuli ^8,10,11,14,16^ while in the primary auditory cortex (A1), locomotion diminishes the depolarization induced by auditory tones ^13,16^. Whether such differences are linked to variable sensitivity of inhibitory circuits to behavioral state transitions across sensory cortical areas remains unexplored.

The control of disinhibitory circuits by active behavioral states has been the focus of considerable scrutiny in V1 ^8,9,15,17–19^. Locomotion, in particular, has been found to influence the activity of VIP- and SST-INs in a context-dependent manner, with the modulation of SST-INs being contingent on the presence or absence of ambient light ^18,19^. In S1, behavioral periods associated with increased whisking (in stationary animals) have been reported to increase L2/3 VIP-INs activity with concomitant decrease in L2/3 SST-INs firing in agreement with the presence of disinhibitory circuits ^5–7,20,21^. Nevertheless, in S1, and in strong contrast to V1, our knowledge regarding the impact of locomotion on the operation of SST- and VIP-INs remains limited. This becomes especially relevant considering that, during navigation, mice typically employ locomotion in conjunction with whisking to explore their surroundings.

Moreover, the mechanisms governing the recruitment of cortical INs, particularly SST-INs, during active behaviors remain unclear. A recent study revealed that modulation of INs activity in V1 by brain states can be largely predicted by the first transcriptomic principal component^17^ . This transcriptomic component suggests that the heightened activity of SST-INs observed in alert states is linked to cholinergic signaling^17^. However, this finding contradicts a prior study that reported a loss of cholinergic modulation in SST-INs in V1 during development^22^. In contrast, experimental evidence does indicate that VIP-INs, which exert inhibition onto SST-INs, rely on cholinergic receptors to positively regulate their in vivo activity in response to transitions in behavioral states^5,15^. Overall, these hypotheses point to neuromodulatory inputs, in particular cholinergic axonal projections to be at the origin of the sensitivity of VIP- and SST-INs to behavioral state transitions. Yet, recent results highlighted that thalamic activity is essential for the large and sustained depolarizations associated with arousal/movement-related signals in L2/3 PNs in V1^23^. These results suggest that behavioral states can modulate neocortical activity not only through neuromodulatory projections such as cholinergic and noradrenergic pathways, but also via glutamatergic thalamic inputs. As SST-INs receive pronounced inputs from local L2/3 PNs it is therefore possible that feedforward activity transmitted via the thalamus participates in the behavioral-dependent modulation of SST-INs in primary sensory regions. Nevertheless, how both neuromodulatory and thalamic signals contribute to the regulation of SST-IN activity during spontaneous behaviors remains uncertain.

Here we used intravital two-photon calcium imaging in S1 and V1 of awake, head-fixed mice, to investigate how transitions in behavioral states (namely whisking and locomotion) modulate the activity of SST- and VIP-INs. We found that in darkness, the recruitment of SST-INs by locomotion was significantly higher in S1 than V1. We also observed that the increase of SST-INs activity in S1 occurred despite the strong recruitment of VIP-INs, a known inhibitor of SST-INs. In addition, neither the strength of inhibition provided by VIP-INs nor the sensitivity of SST-INs to cholinergic signaling was different across sensory regions. Importantly, inactivation of thalamic function using muscimol microinjections limited the increase in activity of SST-INs by locomotion. These results reveal that behavioral state-transitions, associated with changes in arousal, induce sensory modality specific changes to the activity of inhibitory circuits, most likely through inputs coming from the sensory thalamus. In a spiking network model combining a disinhibitory and a thalamo-cortical circuit, we show that various levels of sensory and neuromodulatory drive could indeed explain the various modulations observed in SST-INs activity across conditions. By integrating feedforward thalamic activity transmitted indirectly via L2/3 PNs as well as neuromodulatory information coming from VIP-mediated inhibition and cholinergic innervation, SST-INs occupy a pivotal position in cortical networks by adapting sensory processing to behavioral states in a sensory modality-specific manner.

## Results

### Activity of L2/3 SST-INs in S1 and V1 during spontaneous behavior

We initially focused our attention on the vivo response of SST-INs to behavioral state transitions in the mouse S1. In fact, multiple studies have investigated the activity of disinhibitory circuits during spontaneous behaviors in V1. However, it remains unclear whether inhibitory circuits display similar or variable sensitivity to behavioral state transitions in other sensory cortical areas such as S1. To monitor alterations in firing dynamics of SST-INs we expressed the genetically encoded calcium indicator GCaMP6s selectively in SST-IN using transgenic mice (SST-cre) (**Figure 1A**) and changes in somatic fluorescence of GCaMP6s were used as a proxy for spiking activity. Fluorescence signals were recorded using two-photon imaging in awake head fixed adult mice (>P60) that spontaneously transitioned between rest and locomoting periods on a circular treadmill (**Figure 1A**). Whisking activity was simultaneously acquired using a video camera focused on the mouse snout (**Figure 1A**). This experimental setup allowed us to study how SST-INs activity varied across active behavioral states namely during whisking with and without associated running.

**Figure 1:**
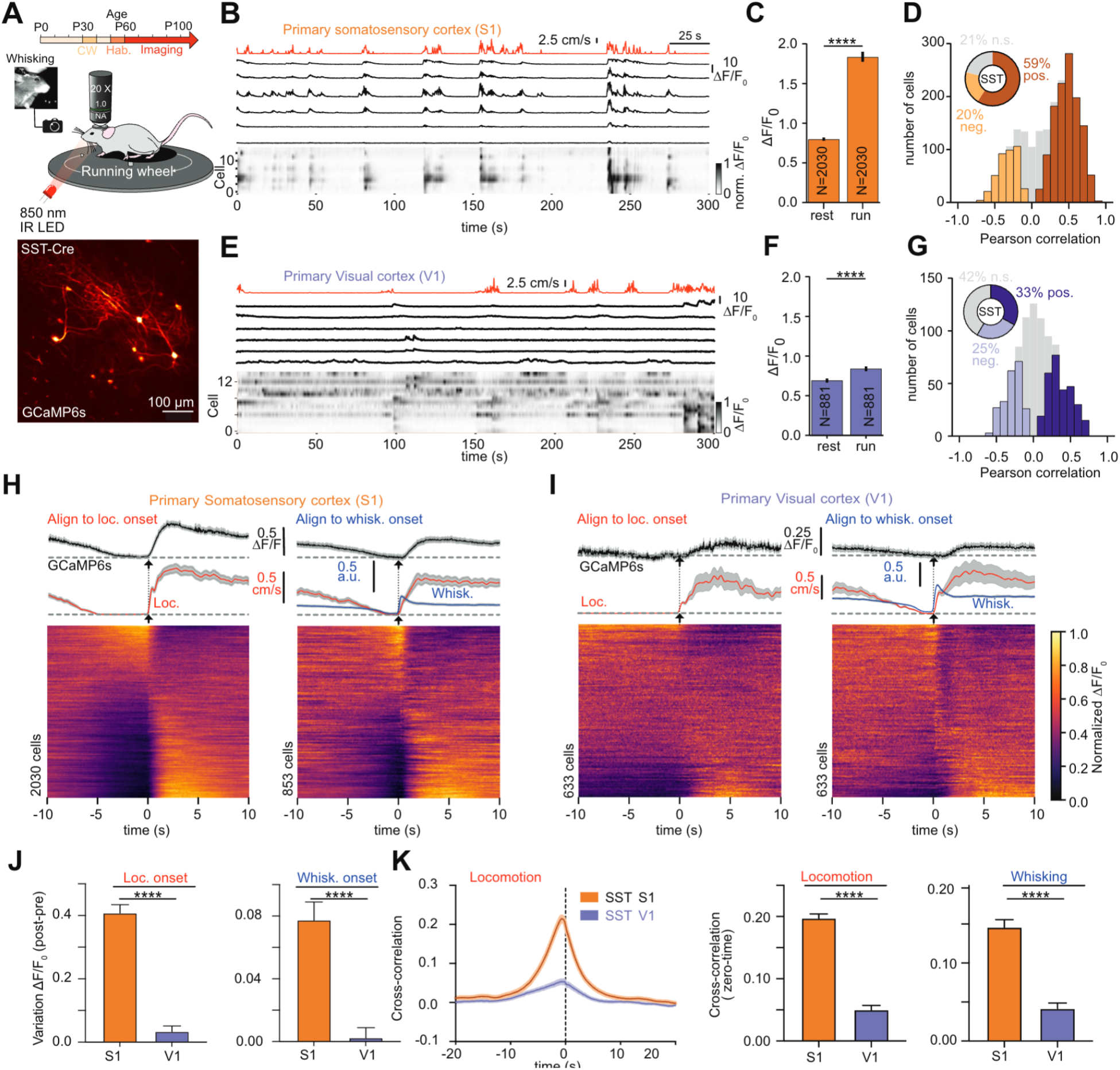
SST-INs in V1 and S1 are differentially modulated by behavioral state transitions. (A) Two-photon (2P) calcium imaging in S1 of awake mice. Mice were first implanted with cranial windows (CW) between P30-P40, and after 10 days of recovery were used for imaging. Recordings started after a 10 days period where mice were habituated to be head fixed and free to run on a circular treadmill. Whisking activity was tracked via a video camera detecting infra-red light focused on mouse snout illuminated with 850 nm LED. Bottom: Two-photon laser scanning microscopy (2PLSM) image (maximum-intensity projection) of SST-Cre positive neurons labelled with GCaMP6s in S1. Scale bar is 100 *μ*m (B) Representative GCaMP6s fluorescence traces (black) shown as ΔF/F0 (up) as well as normalized ΔF/F0 (bottom) for SST-INs displayed in A. Each row represents a neuron sorted by weight on the first principal component (PC) of their activity. Locomotion trace is shown on top (red). (C) Mean ΔF/F_0_ during periods of rest and run (ΔF/F0 rest = 0.80 ± 0.02; ΔF/F0 run = 1.84 ± 0.06; n = 2030 cells in 16 mice, p<0.0001, Mann-Whitney U test). (D) Summary plot of Pearson correlation coefficients between calcium activity and running speed. SST-INs showing statistically significant (see Methods) modulation by locomotion are represented by colored bars, with light orange indicating negative modulation and positive modulation denoted by brown. The pie chart provides an overview of the overall percentage of SST-INs showing positive (dark orange), negative (light orange), or non-significant (grey) correlations with locomotion. (E) Same as B but for SST-INs in the primary visual cortex. (F, G) Same as C and D but for V1. ΔF/F_0_ rest = 0.70 ± 0.03; ΔF/F_0_ run = 0.84 ± 0.03; n = 881 cells in 10 mice, p<0.0001, Mann-Whitney U. (H) Event-triggered average traces of SST-INs GCaMP6s signal in S1 aligned to the start of running (left) or whisking (right) along with running speed (red) and whisking (blue). Normalized ΔF/F is displayed for all analyzed cells (bottom). Traces represent mean ± SEM values across all recorded cells. (I) Same as G but for V1. (J) Difference in ΔF/F following locomotion (left) and whisking (right) events. Values were obtained by averaging ΔF/F trace for each individual SST-INs in a 2 s window before (“pre”) and after (“post”) event onset (see Methods). ΔF/F_Loco S1_ = 0.40 ± 0.03, n = 2030; ΔF/F0_Loco V1_ = 0.03 ± 0.02, n= 633, p<0.0001, Mann-Whitney U; ΔF/F_Whisk S1_ = 0.08 ± 0.01, n = 853, n = 2030, ΔF/F_Whisk V1_ = 0.002 ± 0.007; n = 853 cells in 10 mice, p<0.0001, Mann-Whitney U. (K) Cross-correlation between in vivo calcium activity and running speed for SST-INs in S1 and V1 (left). (right) Average correlation coefficient values at zero-time for locomotion and whisking for SST-INs recorded in V1 and S1 (*ρ*_loco_ S1 = 0.22 ± 0.01, n = 2030, *ρ*_loco_ V1 = 0.06 ± 0.01, n = 883, p<0.0001, Mann-Whitney U; *ρ*_whisk_ S1 = 0.15 ± 0.01, n = 2030, *ρ*_loco_ V1 = 0.05 ± 0.01, n = 883, p<0.0001, Mann-Whitney U).

In our initial analysis, we categorized recording periods based on whether animals were engaged in whisking, locomotion, or neither. Periods during which whisking or locomotion were not detected were classified as resting (see Star Methods). Through the quantification of average GCaMP6s fluorescence levels we found that in S1, SST-INs displayed markedly higher activity levels during periods of locomotion compared to resting (**Figure 1A-C**). This engagement of SST-INs during locomotion in S1 was also observed when we investigated the response at the single cell level. We quantified the correlation between baseline activity and running speed for individual SST-INs and we observed a predominantly positive correlation coefficient across SST-INs in S1 (**Figure 1D**) with the majority (59%) of SST-INs exhibiting a statistically significant positive modulation in response to locomotion (**Figure 1D**).

These results contrasted with previous reports in V1, where in darkness, locomotion had on average, no impact on SST-INs activity^18,19^. To test if this functional difference was linked to our experimental conditions or reflected a previously unknown region-specific modulation of SST-INs by locomotion, we repeated the experiments but now monitoring SST-INs activity in V1 (**Figure 1E**). In agreement with previous studies we observed that the modulation of SST-INs by locomotion in the dark in V1 was quite modest^18,19^ (**Figure 1E,F**). This difference in activity of SST-INs during locomotion between S1 and V1 was also observed at the single cell level. In contrast to S1, our observations in V1 revealed a comparable ratio of SST-INs exhibiting positive (33%) and negative (25%) correlations between running speed and spontaneous activity (**Figure 1G**).

In order to better estimate the rate of change in calcium levels in SST-INs between S1 and V1 during active behaviors we generated population averages of all running bouts aligned to locomotion onset (**Figure 1H**). In these population averages it was possible to observe that locomotion induced changes in ΔF/F_0_ in SST-INs that were significantly larger in S1 than in V1 (**Figure 1J, I**). Similar findings were obtained when we monitored activity of SST-INs associated with whisking periods (**Figure 1J**). The differential sensitivity of SST-INs between the two sensory areas to behavioral state transitions was particularly evident when comparing the cross-correlation curves between spontaneous activity of SST-INs and the locomotion or whisking traces (**Figure 1K**). Altogether these results suggest that in the neocortex, behavior-dependent modulation of SST-INs activity is not uniform across sensory regions.

### Contribution of whisking and locomotion to SST-INs modulation in S1

Our results indicate that in S1 active behaviors like whisking and locomotion, result in increased activity of SST-INs. However, locomotion in mice is closely linked to the motion of their whiskers (**Figure 2A**, ^24^). In contrast, whisking can take place in both run and rest conditions (**Figure 2A**; labelled as “Loc. +Whisk.” and “Whisk.-only” respectively). We thus decided to further refine how those two different whisking states impacted the activity of SST-INs. In our experimental conditions mice spend an average 29 % of their time whisking with associated locomotion and 15% of their time in whisking only periods (**Figure 2A-B, Figure S1C** and Star Methods). Activity of SST-INs was quite distinct between the two whisking states. Specifically, SST-INs were positively modulated during whisking periods with associated locomotion. Yet, no modulation or a tendency to inhibition was observed in whisking only (no locomotion) segments in agreement with previous works^5–7,20^ (**Figure 2C-D**). To further quantify how the different behavioral states influenced SST-INs activity we separated calcium activity, and we defined whisking only modulation index (WOMI) and whisking modulation index (WMI) similar to our previous LMI calculation (see Star Methods). We observed that both WMI and LMI were clearly positive and rather similar in agreement with the observation that in our experimental conditions most whisking bouts were associated with locomotion. In contrast WOMI, was only slightly positive with a large percentage of cells displaying negative modulation (**Figure 2E**).

**Figure 2:**
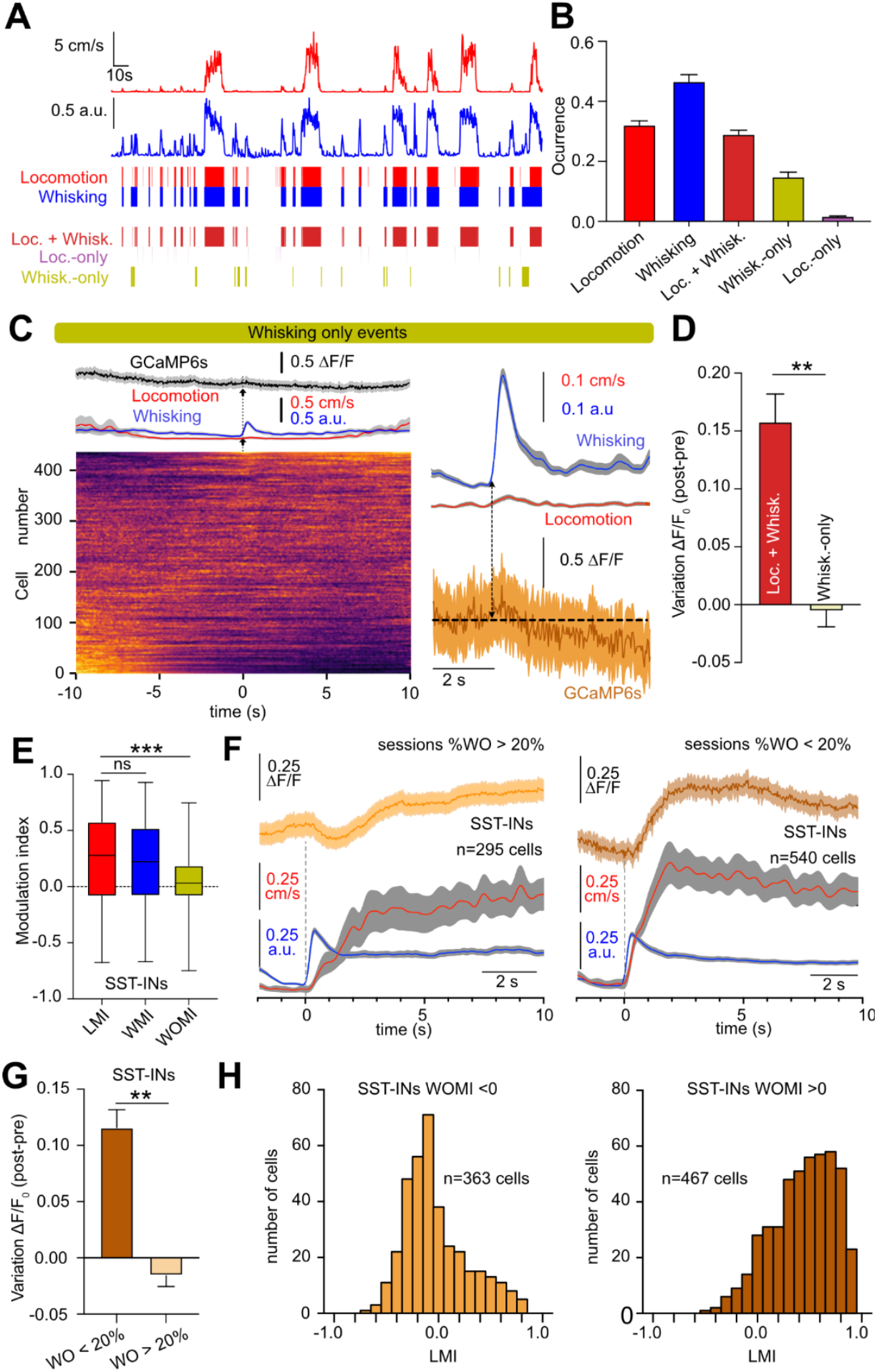
Recruitment of SST-INs in S1 only occurs with behavioral state transitions characterized by increased locomotor activity. (A) Example session illustrating simultaneous recording traces for whisking activity (blue) and running speed (red). Those traces were used to define “Locomotion” and “Whisking” periods, then separated into whisking-only (“Whisk.-only”), locomotion-only (“Loc.-only”) and locomotion and whisking (“Loc.+Whisk.”) events (see Star Methods). (B) Fraction of time of the different behavioral events during the imaging sessions (N= 85). (C) (left) Event-triggered average traces of SST-INs GCaMP6s fluorescence signal in S1 aligned to whisking onset. Only whisking bouts not associated with locomotion were used (whisking only). Normalized ΔF/F is displayed for all analyzed cells (bottom). Traces represent mean ± SEM values across all recorded cells. (right) Enlarged view of average traces shown on the left. GCaMP6s trace is average of all cells. (D) Variations in ΔF/F following whisking events shown in C, calculated as the difference between “pre” and “post” event ΔF/ F values by averaging a 2 s window before and after event onset respectively (see Methods). Note that during whisking only, there is either no effect or a slight tendency to inhibit SST-INs. ΔF/F_0_ Loc.+Whisk. = 0.16 ± 0.02, n = 438; ΔF/F_0_ Whisk only = -0.01 ± 0.01; n = 436 cells in 10 mice, p<0.001, Mann-Whitney U. (E) Summary plot of average mean locomotion, whisking and whisking only modulation indices (LMI = 0.26 ± 0.01, n=2030; WMI = 0.21 ± 0.01, n=856; WOMI = 0.06 ± 0.01, n=830) (F) Event-triggered average traces of SST-INs GCaMP6s signal (ΔF/F) in S1 aligned to whisking onset. Results were separated according to percentage time mice spent whisking only during the imaging session. Average of GCaMP6s fluorescence from sessions with whisking only periods longer than 20% of the recording time is shown on the left and less than 20% is illustrated on the right. Traces represent mean ± SEM values across all recorded cells. (G) Same as D but for events displayed in panel F, (ΔF/F_0_ Whisk<20% = 0.11 ± 0.02, n = 602; ΔF/F_0_ Whisk<20% = -0.02 ± 0.01, n = 251, p<0.0001, Mann-Whitney U). (H) Histograms of the distribution of locomotion modulation indices (LMI = (RL – RS)/(RL + RS), where RL and RS are the mean ΔF/F_0_ during locomotion and stationary periods, respectively), for SST-INs that displayed negative or positive WOMI.

During our imaging sessions, we observed notable variations in occurrence of whisking only periods (% time WO = 14.7 ± 1.7%, CV = 106%, n = 85 sessions). Hypothesizing that a higher proportion of WO activity would provide a better opportunity to sample differences in the response profile of SST-INs between the two whisking states we categorized sessions using a 20% threshold (**Figure 2F, G**). Examining whisking bouts, we observed distinct variations in the response of SST-INs between the two defined groups. Sessions with predominantly whisking-only activity showed a decrease in the activity of SST-INs post-whisking onset, while sessions with minimal whisking without locomotion exhibited an increase in SST-INs activity (**Figure 2F,G**). Overall, these results indicate that in S1, whisking at rest induces a decrease in SST-INs activity, in agreement with previous reports ^6,7,21^. On the other hand, when mice engage in active behavioral states involving locomotion, activity of SST-INs is increased, specifically in S1, but not in V1 (**Figure 1**). These findings indicate that distinct active behavioral states exert varying effects on the activity of S1 SST-INs.

Morphoelectric and transcriptomic analysis have highlighted that SST-INs, even within L2/3, are a highly heterogeneous neuronal population^25,26^. We probed if sensitivity to locomotion was also heterogeneous across individual SST-INs in S1. To evaluate this, we separated SST-INs according to their response to WOMI. We observed that SST-INs that displayed a negative WOMI, were on average less sensitive to locomotion (**Figure 2H**). In contrast, SST-INs with positive WOMI mostly increase their activity with locomotion (**Figure 2H**). Variable modulation of SST-INs for locomotion was observed even within single fields of view, suggesting that variability in SST-IN responses cannot be attributed to distinct behavioral states (**Figure S1**). In summary, these findings suggest that SST-INs in L2/3 exhibit varying degrees of sensitivity to transitions in behavioral states which might be linked to the presence of various subtypes of SST-INs.

### VIP-INs are equally engaged by locomotion in V1 and S1

Having observed that in S1, SST-INs are recruited during active behavioral states, specifically during locomotion, we proceeded to investigate the underlying neuronal mechanisms involved in such modulation. Taking into account previous works we identified three main potential pathways that could participate in controlling L2/3 SST-INs activity during spontaneous behaviors: (i) inhibitory top-down modulation via VIP-INs^15,20,27,28^ (ii) increased local recurrent excitatory L2/3 PNs activity^19,23^ and (iii) direct excitatory neuromodulation (e.g. via direct cholinergic inputs)^22^. We initially focused on VIP-INs, which are known to be important inhibitors of SST-IN function ^4,20,27,28^ and are strong recipients of neuromodulatory inputs ( *e*.*g*. cholinergic) associated with active behavioral states^5,29^. Therefore, we studied the contribution of VIP-INs to the activity patterns of SST-INs during behavioral state transitions. We also explored the possibility that differential activity/function of VIP-INs could be at the origin of the distinct sensitivity of SST-INs in V1 and S1 to locomotion.

We started by examining how activity of VIP-INs varied in S1 during the same behavioral transitions as we described for SST-INs (**Figure 3A-C**). In line with previous findings^3,4^, we observed an increase in activity of VIP-INs during ambulatory periods (**Figure 3A, B, D**) with the majority of VIP-INs displaying a positive modulation by locomotion (**Figure 3E**). A similar observation was obtained while estimating Pearson correlation between GCaMP6f or GCaMP6s fluorescence and running speed. VIP-INs displayed on average a positive correlation (**Figure 3F**) with only a very small fraction (6%) of VIP-INs displaying a statistically significant decrease in activity (**Figure 3F**) during locomotion. Despite the similar average positive modulation by locomotion between VIP- and SST-INs in S1, we observe that at the population level, VIP-INs displayed a higher sensitivity to behavioral state transitions than SST-INs (**Figure S2**). This was reflected into a higher correlation coefficient between spontaneous activity and running speed (**Figure S2A**) and different response profile between running speed and the variation in ΔF/F_0_ when compared with SST-INs (**Figure S2B**). Similar findings were observed when analyzing whisking and whisking only periods (**Figure S2C-F**). Both whisking and whisking only resulted in increased activity of VIP-INs, which (**Figure S2C-F**) contrasted SST-INs, where whisking only did not induce any detectable increase in activity (**Figure 2D,E**). Such increased sensitivity of VIP-INs to active states is probably linked to the enrichment of VIP-INs in nicotinic cholinergic receptors when compared to SST-INs (**Figure S2G, H** and ^5^). Importantly, and in contrast to SST-INs we observed comparable modulation of VIP-INs by locomotion in V1 and S1 (**Figure 3C, G-I**) ^19^. Analogous findings were obtained while studying whisking periods.

**Figure 3:**
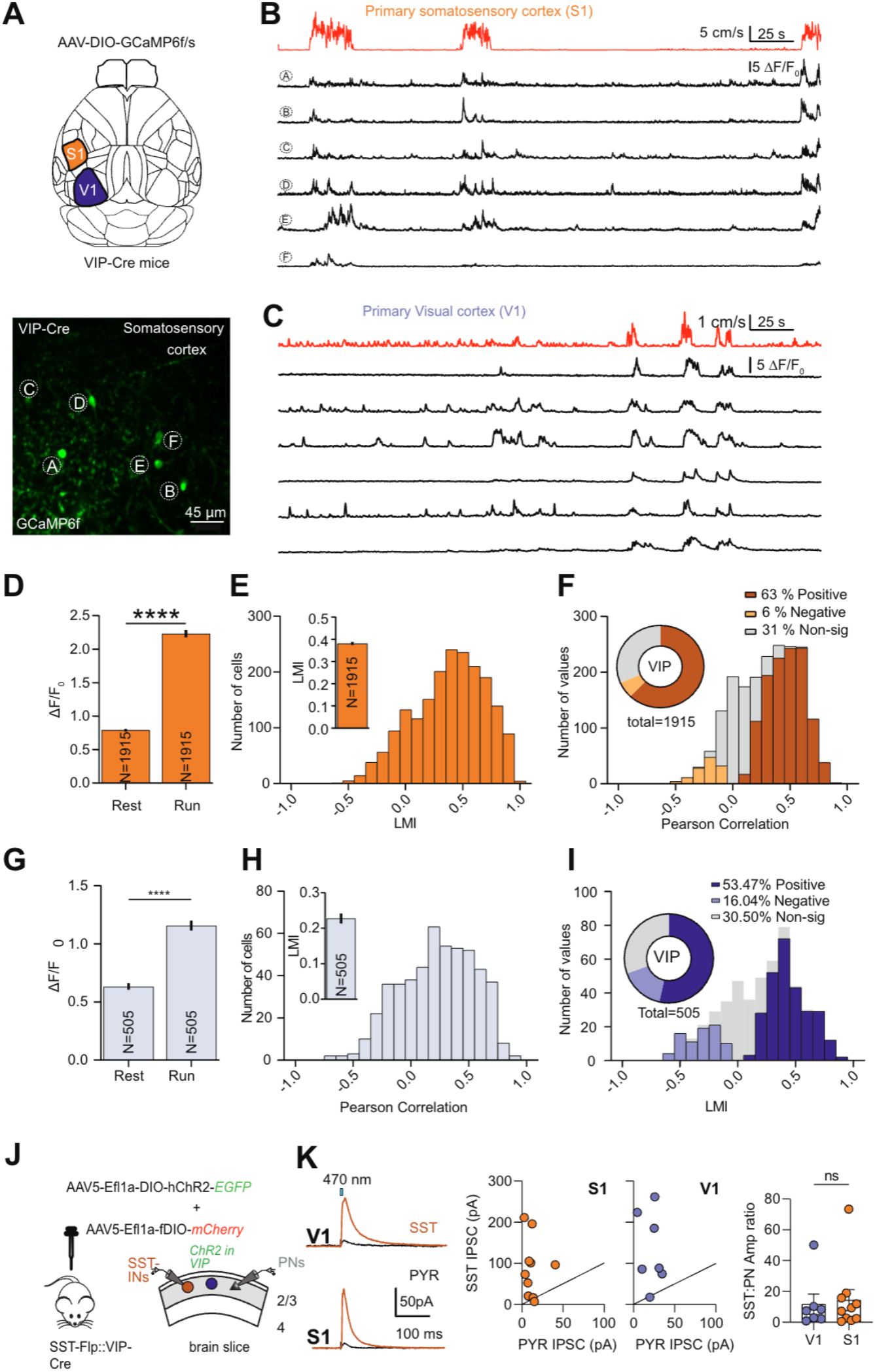
VIP interneurons are strongly modulated by locomotion in S1 and V1. A) (top) Illustration of a mouse brain depicting the location of S1 and V1. (bottom) 2PLSM image (maximum-intensity projection, MIP) of VIP-Cre positive neurons labeled with GCaMP6f in S1. B) Representative traces of in vivo calcium (GCaMP6f) activity in VIP-INs. Locomotion trace is shown on top (red) followed by the calcium activity (green). C) Same as B but for VIP-INs in V1. D) Mean ΔF/ F_0_ during periods of rest and run, (ΔF/F0 rest = 0.80 ± 0.02; ΔF/F0 run = 2.23 ± 0.05; n = 1915 cells in 11 mice, p<0.0001, Mann-Whitney U test). E) Histogram of locomotion modulation indices for VIP-INs in S1. Inset depicts the average LMI value (LMI = 0.38 ± 0.01, n= 1915). F) Distribution of Pearson correlation coefficients between calcium activity and locomotion trace for VIP-INs in S1. Colored bars indicate VIP-INs that are statistically significantly modulated by locomotion. The pie chart shows the overall percentage of VIP-INs showing positive, negative, or non-significant correlations with locomotion. G, H, I) Same as D, E, F but for VIP-INs in V1. J) Illustration of the double transgenic mouse line used for targeted protein expression in VIP- and SST-INs respectively. Adeno-associated virus (AAV) coding Cre-dependent expression of Channelrhodopsin2-EGFP along with a second AAV coding FLP-dependent expression of mCherry were infected in primary visual (V1) and primary sensory (S1) cortices of VIP-Cre::SST-Flp mice. Schematic diagram of experimental approach where VIP-INs were photo-activated via ChR2 while a PN and neighboring SST-IN were patched in L2/3. K ) ( left ) Example traces of IPSCs simultaneously recorded in PNs and SST in V1 and S. (middle) Scatter plot of IPSCs amplitudes recorded in SST-INs and PNs in S1 and V1 following VIP-INs photostimulation. (right) Ratio of light-evoked IPSCs in V1 and S1 (V1: n = 7 cells, 2 mice ; S1: n = 10 cells, 2 mice). Mann Whitney test p=0.8125. Error bars represent SEM.

We further investigated a potential difference in the strength of inhibition from VIP-INs to SST-INs between the two primary sensory regions using ex-vivo whole-cell patch-clamp recording. For that we used animals that expressed the Cre Recombinase in VIP-INs while the FLP recombinase was expressed in SST-INs (**Figure 3J**). Such transgenic mice allowed us to selectively express channelrhodopsin (ChR2) in VIP-INs and to record light-evoked IPSCs from fluorescently labeled SST-INs (**Figure 3J**). We normalized responses through comparison to simultaneously patched neighboring PNs. As previously reported, we observed that VIP-mediated inhibition was higher in SST-INs when compared to local PNs, but that this ratio was similar between S1 and V1 (**Figure 3K**). Taken together, our findings indicate that changes in VIP-IN-mediated inhibition are likely not responsible for the differential impact of locomotion and whisking in SST-INs between V1 and S1.

### VIP-INs inhibit SST-INs during active states in S1

According to the classical “disinhibitory” scenario^27^, the activation of VIP-INs is expected to decrease the activity of SST-INs. Our observation that during rest/run transitions both VIP- and SST-INs in S1 increase their activity is not easy to reconcile with such a hypothesis. Alternatively, theoretical work suggested that VIP-INs activation in certain conditions can result in unexpected increase in SST-INs function through disinhibition of local PNs^30^. Hence, we performed experiments to assess the influence of manipulating VIP-INs activity on the recruitment of SST-INs during active behaviors in S1.

To achieve this, we used VIP-Cre::SST-Flp transgenic mice to express the inhibitory DREADD hM4Di in VIP-INs while GCaMP6s was selectively expressed in SST-INs (**Figure 4A-B**). Mice were initially imaged under control conditions, and then re-imaged 30 minutes after the injection of CNO (1-10mg/Kg) to monitor potential changes in the activity of SST-INs (**Figure 4B**). In both control conditions and after CNO injection, we observed an increase in average activity of SST-INs during locomotion (**Figure 4C**). Interestingly, when investigating the modulation at the single cell level, we found that SST-INs exhibited greater sensitivity to locomotion after CNO treatment, as evidenced by elevated LMI and Pearson correlation values (**Figure 4D**). The proportion of negatively modulated cells was also reduced in CNO treated mice (**Figure 4D**). We subsequently averaged activity over all whisking periods aligning for whisking onset (**Figure 4E**). In control condition we observed that immediately after whisking there was a decrease in SST-INs activity that progressively increased as locomotion speed increased (**Figure 4E-F**). In CNO treated conditions, the inhibitory effect after whisking was absent and overall, there was a larger increase in positive modulation of SST-INs by locomotion (**Figure 4F-E**). Detailed analysis in the variation of SST-INs activity with running speed also revealed that after CNO treatment a similar speed produced a larger increase in ΔF/F_0_ signal in SST-INs (**Figure 4H, I**). These results suggest that during active states VIP-INs have an overall inhibitory action in SST-INs and raises the question about the underlying mechanism responsible for the active recruitment of L2/3 SST-IN during locomotor activity.

**Figure 4:**
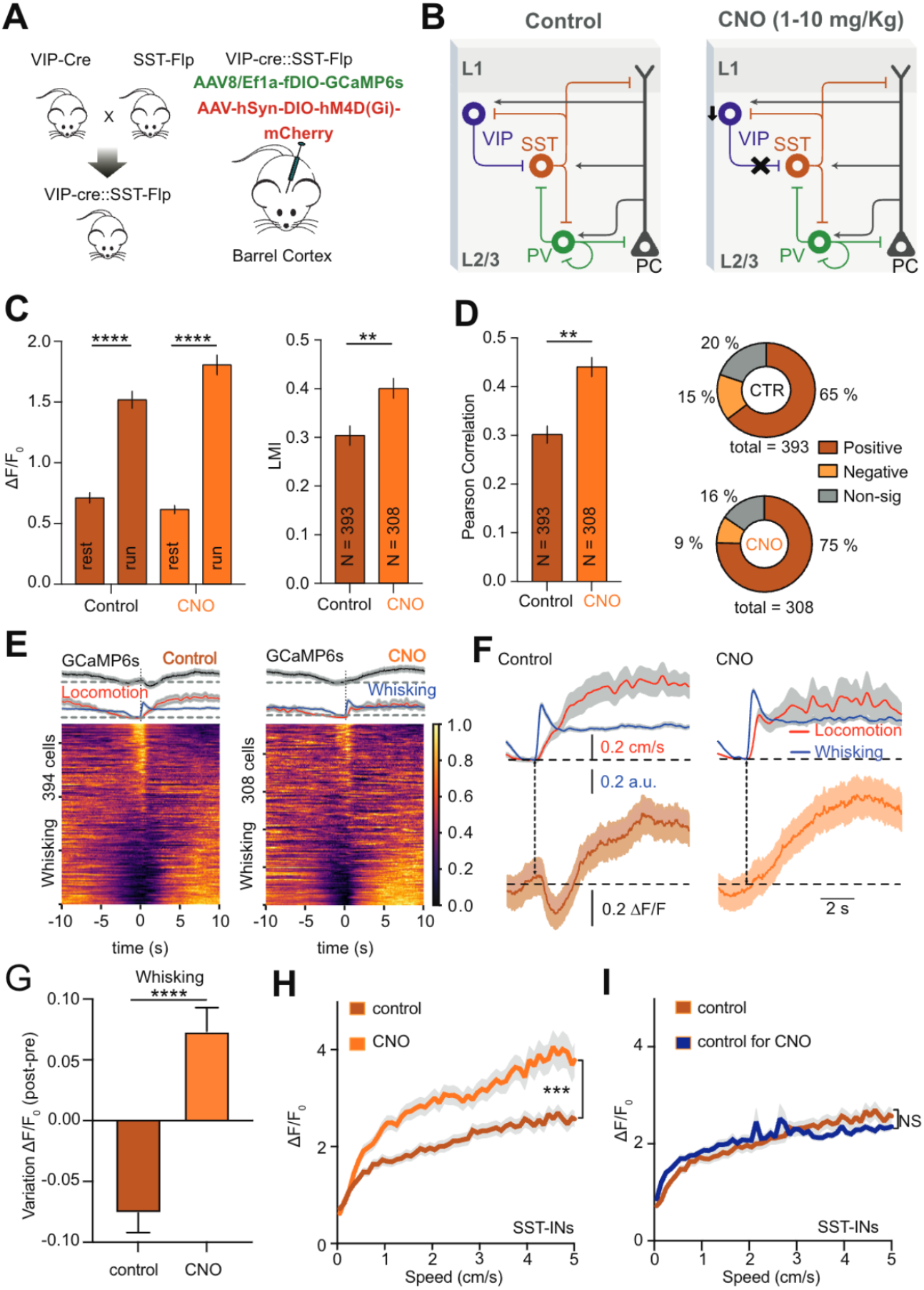
VIP-INs mainly inhibit SST-INs during spontaneous behaviors in S1. A) Simplified illustration of the double transgenic mouse line used to target protein expression in VIP- and SST-INs respectively based on the Cre/Lox and FLP/FRT system. Adeno-associated viruses (AAV) were used to express in a Cre-dependent manner Gi-coupled inhibitory (hM4Di) designer receptors exclusively activated by designer drugs (DREADD) in VIP-INs. A second AAV was used to drive in a FLP-dependent manner the expression of GCaMP6s in SST-INs. B) Schematic illustration of the canonical cellular organization in S1 and the expected negative modulation of VIP-INs via activation of hM4Di by CNO injection. C) Mean ΔF/F0 during periods of rest and run in control conditions and 30 after CNO injection (left), (ΔF/F_0_ rest, control = 0.71 ± 0.02; ΔF/F_0_ run, control = 1.5 ± 0.07; n = 393 cells in 5 mice, p<0.0001, Mann-Whitney U test; ΔF/F_0_ rest, CNO = 0.61 ± 0.04; ΔF/F0 run, CNO = 1.81 ± 0.08; n = 308 cells in 5 mice, p<0.0001, Mann-Whitney U test). Summary plot of LMI values obtained for SST-INs in control and after CNO injection (right), (LMI control = 0.30 ± 0.02, n = 393; LMI CNO = 0.4 ± 0.02, n = 308; p<0.001, Mann-Whitney U test). D) Variation in Pearson correlation values between running speed and GCaMP6s fluorescence signals for SST-INs recorded in control and CNO treated mice (ρ_loco_ control = 0.30 ± 0.02, n = 393, ρ_loco_ CNO = 0.44 ± 0.02, n = 308, p<0.0001, Mann-Whitney U test). Percentage of SST-INs exhibiting statistically significant positive, negative, or non-significant Pearson correlation values is shown on the right. E) Event-triggered average traces of SST-Ins GCaMP6s signals in S1 aligned to the onset of whisking along with running speed (red) and whisking (blue). Normalized ΔF/ F is displayed for all analyzed cells (bottom). Results are displayed for control conditions (left) and after CNO injection (right). Traces represent mean ± SEM values across all recorded cells. F) Enlarged view of average traces shown in E. G) Variations in ΔF/F following whisking events shown in F, calculated as the difference between “pre” and “post” event ΔF/F values by averaging a 2 s window before and after event onset respectively (see Methods). Values were calculated for each individual SST-INs. Note that in control conditions whisking induces a brief decrease in GCaMP6s fluorescence that is absent in CNO treated mice. H) Variation in ΔF/F values for SST-INs with running speed for control and CNO conditions. I)Same as H but comparing the variation in ΔF/F values for SST-INs with running speed in control mice used for the CNO experiments and in the mice used in the rest of the study. It is important to note that the two curves exhibit no statistical differences, suggesting consistent variation of ΔF/F with speed across different batches of animals.

### Visual stimulation alters SST-INs activity during active states in V1 but not in S1

The variability in VIP-dependent disinhibitory circuits is unlikely to be the source of the positive modulation of SST-INs during active states and to be at the origin of the observed differences in SST-IN activity between S1 and V1. We also ruled out potential differences in the expression of cholinergic receptors in SST-INs between the two sensory regions^22^ (**Figure S3**). We thus looked for alternative pathways that could explain the positive modulation of SST-INs during active states in S1. Recent work revealed that thalamic activity is essential for the arousal/movement-related activation of PNs in V1^23^. Inspired by those findings we tested if feedforward signals transmitted to the cortex were at the origin of the observed modulation of SST-INs by active behavioral states.

In V1, SST-INs display modest to no modulation by locomotion in the dark^4,18,19^ but show increased activity during active states when mice are facing a grey screen^18,19^. These results are compatible with the idea that increased activity of thalamic neurons contribute to defining the magnitude or direction of modulation of cortical SST-INs to behavioral transitions in a sensory modality specific manner. We thus investigated whether in our working conditions visual stimulation which is expected to increase the activity of the feedforward pathways into V1 altered the sensitivity of SST-INs to behavioral transitions from rest to run in both S1 and V1. To achieve this, we monitored the activity of SST-INs in both V1 and S1 under two conditions: in the dark and in front of a grey screen (**Figure 5A**). We assessed changes in activity for both VIP- and SST-INs from rest/run transitions by measuring variations of LMI as well as the Pearson correlation between spontaneous activity and running speed (**Figure 5**). Our observations revealed that VIP-INs activity remains relatively constant between S1 and V1, regardless of the dark or grey screen conditions (**Figure 5A-D**). In contrast, SST-INs in V1 displayed a significantly greater sensitivity to locomotion when facing a grey screen. Interestingly, the modulation of SST-INs in S1 by locomotion appeared to be relatively unaffected by the dark versus grey screen conditions (**Figure 5A-D**). These findings indicate that visual stimulation plays a crucial role in enhancing the sensitivity of SST-INs in V1 to changes in locomotion as previously reported^18,19^. However, this effect seems limited to V1 and is not observed in S1. This led us to investigate whether whisking signals would be necessary for the recruitment of SST-INs in S1 during locomotion, potentially operating in a manner analogous to the influence of visual stimulation in V1. To test that we compared the activity of SST-INs in control mice and in mice where whiskers were trimmed. Notably, the activity of SST-INs during locomotion was significantly diminished by whisker trimming on day 0 and day 3 post-trimming (**Figure S4**). Upon careful examination, we observed that there was no tactile contact with the mouse whiskers during mouse running in our experimental setup. This suggests that the most probable source of the enhanced activity in SST-INs during locomotion is thus the generation of reference signals in the whisker follicle or whisker pad derived from whisker movements in the air (see discussion and ^31^). Altogether these observation suggest that activity of feedforward pathways is important for sensitivity of SST-INs to behavioral state transitions.

**Figure 5:**
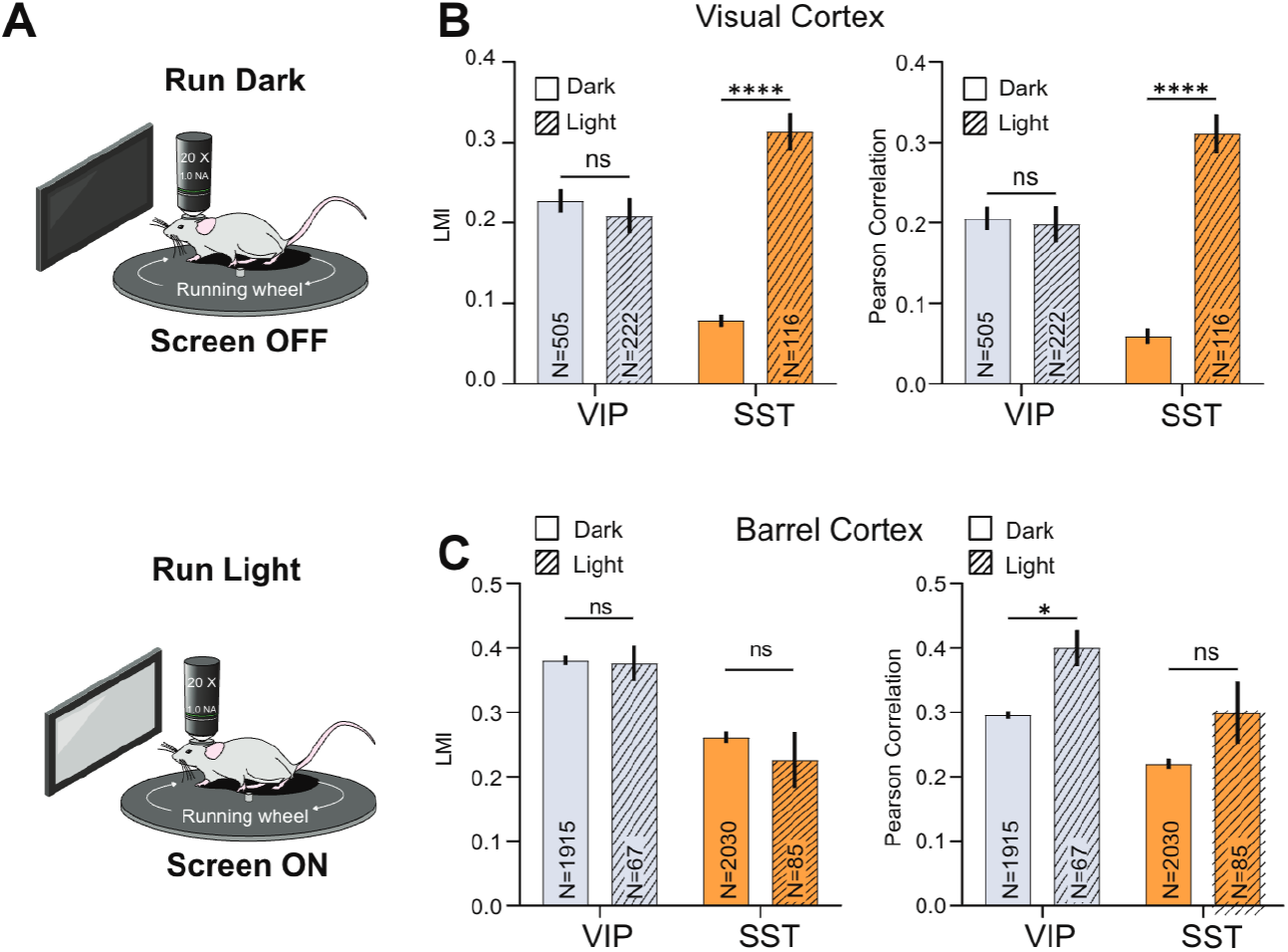
Visual stimulation modulates the recruitment of SST-INs by locomotion in V1 but not in S1. A) Schematic of the experimental setup used to probe the effect of visual stimulation in the behavioural state-dependent modulation of the activity of VIP and SST-INs in V1 and S1. B) Summary plots of LMI and Pearson correlation between running speed and GCaMP6 fluorescence signal for VIP- and SST-INs in the dark and facing a grey screen. LMI = 0.23 ± 0.01, n =505, for VIP in the dark versus 0.21 ± 0.02, n = 222, in light; p > 0.9999, and 0.08 ± 0.01, n = 881 for SST in the dark versus 0.31 ± 0.02, n = 116 in light, p < 0.0001. Pearson Correlation: 0.21 ± 0.01, n = 505, for VIP in the dark versus 0.20 ± 0.02, n = 222, in light; 0.06 ± 0.01, n = 881, for SST in the dark versus versus 0.31 ± 0.02, n = 116 for SST-INs in light, p < 0.0001, Mann-Whitney U test. C) Same as B but for S1. Values represent mean ± SEM. LMI = 0.34 ± 0.02, n=194, for VIP in the dark versus 0.37±0.03, n=67, in light; p>0.05, and 0.24±0.02, n=367 for SST in the dark versus 0.23 ± 0.04, n = 85 in light, p > 0.05. Pearson Correlation: 0.30 ± 0.01, n = 194, for VIP in the dark versus 0.40 ± 0.03, n = 67, in light, p < 0.01, Mann-Whitney U test; 0.27 ± 0.02, n = 367, for SST in the dark versus 0.30 ± 0.05, n = 85 for SST-INs in light, p > 0.05, Mann-Whitney U test. Values represent mean ± SEM.

### Thalamic activity is required for the modulation of SST-INs by spontaneous behaviors

As visual stimuli altered sensitivity of SST-INs to locomotion in V1 but not S1 we reasoned that most likely thalamic activity is required for the modulation of SST-INs by active behavioral states. We raised the hypothesis that in the dark, whisking leads to an increase in the activity of the somatosensory thalamus^32^ that is subsequently relayed to S1 and explains the sensitivity of SST-INs to locomotion in the dark but not in V1. To directly test this hypothesis, we inactivated the somatosensory thalamus (targeted on VPM coordinates, see Star Methods) with microinjections of the GABA(A) agonist muscimol (0.5 mM) and monitored the effect in SST-INs during resting and active states. We first validated that we were able to effectively inactivate the somatosensory thalamus. For that we conducted in vivo calcium imaging experiments in S1 while monitoring activity of PNs as well as SST-INs in response to whisker stimulation in control conditions and after muscimol injections in the somatosensory thalamus (**Figure 6**). We observed that in control conditions whisker stimulation induced a clear increase in the activity of both SST-INs and PNs (**Figure 6A-D**). In contrast, after muscimol injection this increase was absent in both cell types (**Figure 6A-D**). These results confirmed the effectiveness of muscimol in reducing thalamic activity. Following the conclusion of the experiment, we confirmed the accurate targeting of the VPM through muscimol injections (**Figure S5**). On multiple occasions, we observed fluorescence signals in the POM, suggesting that our approach may not effectively distinguish between VPM and POM inactivation (see discussion). Subsequently, we monitored the modulation of SST-INs in S1 by whisking and locomotion in control conditions and after thalamus inactivation (**Figure 6E-F**). Interestingly when we evaluated response of SST-INs to both whisking and locomotion there was a clear modification of the response. In muscimol conditions there was a significant decrease in the response of SST-INs during locomotion. This effect translated into a decreased LMI both at the single cell level as well as when comparing average values across the difference tested mice (**Figure 6G-J** ). The Pearson correlation between spontaneous activity and running speed was also significantly reduced by thalamus inactivation (**Figure 6J**). Plotting variation in ΔF/F vs running speed revealed once more that in control conditions SST-INs activity increased with running speed and that this increase is significantly reduced after muscimol injections (**Figure 6K**). Overall, our results suggest that in S1, modulation of SST-INs by spontaneous behaviors requires thalamic activity.

**Figure 6:**
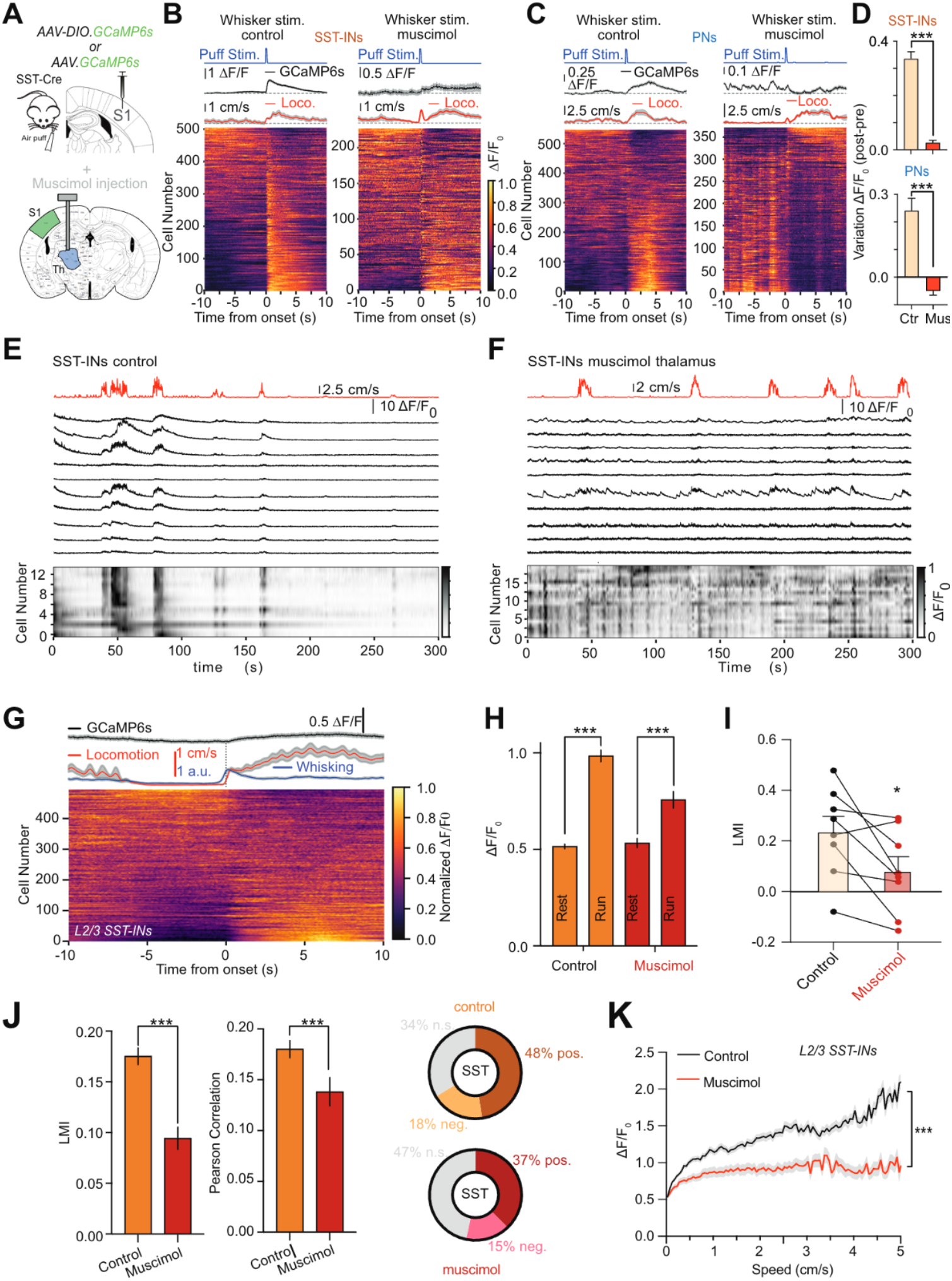
Thalamic activity is required for the modulation of SST-INs by locomotion in S1. A)Schematic illustration of experimental approach used to monitor effectiveness of thalamus inactivation using muscimol injection in somatosensory thalamus . Whisker deflection was achieved using localized air puff while cortical activity in S1 was monitored using in vivo two-photon calcium imaging in both SST-INs and local PNs. B) Event-triggered average traces of SST-INs GCaMP6s signals in S1 aligned to whisker puff along with running speed (red). Normalized ΔF/F is displayed for all analyzed cells (bottom). Results are displayed for control conditions (left) and for mice previously injected with muscimol in the thalamus (right). Traces represent mean ± SEM values across all recorded cells. C) Same as B but for PNs. D) Variations in ΔF/F following puff stimulation events shown in B and C, calculated as the difference between “pre” and “post” event ΔF/F values by averaging a 2 s window before and after event onset respectively (see Methods). E) Representative GCaMP6s fluorescence traces (black) shown as ΔF/F_0_ (up) as well as normalized ΔF/F_0_ (bottom) for SST-IN s in S1 f or mice spontaneously transitioning between rest and active behavior periods. Each row represents a neuron sorted by weight on the first principal component (PC) of their activity. Locomotion trace is shown on top (red). F) Same as A but for mice treated with muscimol injection in the somatosensory thalamus (VPM and POM). G) Event-triggered average traces of SST-INs GCaMP6s signals in S1 aligned to the onset of locomotion along with running speed (red) and whisking (blue). Normalized ΔF/F is displayed for all analyzed cells (bottom). Results are displayed for mice previously injected with muscimol in the thalamus. Traces represent mean ± SEM values across all recorded cells. H) Mean ΔF/F0 during periods of rest and run in control conditions and 30 min microinjection of muscimol in the somatosensory thalamus (ΔF/F_0_ rest control = 0.53 ± 0.01; ΔF/F0 run control = 1.01 ± 0.03; n = 1657 cells in 8 mice, p<0.0001, Mann-Whitney U test; ΔF/F0 rest muscimol = 0.53 ± 0.02; ΔF/F_0_ run muscimol = 0.76 ± 0.04; n = 541 cells in 8 mice, p<0.0001, Mann-Whitney U test). I) Summary plot of average LMI obtained in control and after muscimol injection per the different individual mice tested (LMI_control_ = 0.23 ± 0.06, LMI_muscimol_ = 0.08 ± 0.06, 8 mice, p = 0.0391, Wilcoxon test. J) Summary plots for LMI (left) and Pearson correlation values (middle) between running speed and calcium activity for SST-INs in S1 obtained in control and after muscimol injection in the thalamus (LMI_control_ = 0.18 ± 0.01, n = 1657, LMI_muscimol_ = 0.10 ± 0.01, n = 541, p < 0.0001 Mann-Whitney U test; Pearson, ρ_control_ = 0.18 ± 0.01, n = 1657, ρ_muscimol_ = 0.14 ± 0.01, n = 541, p < 0.05 Mann-Whitney U test) . Percentage of SST-INs exhibiting statistically significant positive, negative, or non-significant Pearson correlation values is shown on the right. K) Variation in ΔF/F values for SST-INs with running speed for control and muscimol injection conditions.

### In S1 locomotion is associated with increased L2/3 pyramidal cell activity

Experimental evidence indicate that SST-INs are not contacted directly by thalamic inputs^33–35^. Yet, SST-INs can be driven by thalamic activity indirectly via their important connectivity with L2/3 PNs^36,37^. To test this, we monitored the in vivo spiking activity of L2/3 in S1 during behavior state transitions namely running and whisking in control conditions and after muscimol injections in the somatosensory thalamus (**Figure 7**). GCaMP6s was expressed in pyramidal neurons using GCaMP6-expressing adeno-associated virus (AAV.hsyn.GCaMP6s, see Star Methods). Due to the markedly higher prevalence of glutamatergic neurons in the neocortex, the use of hsyn, which is pan-neuronal promoter, still results in the majority of GCaMP6s positive cells to be predominantly excitatory neurons^38^.

**Figure 7:**
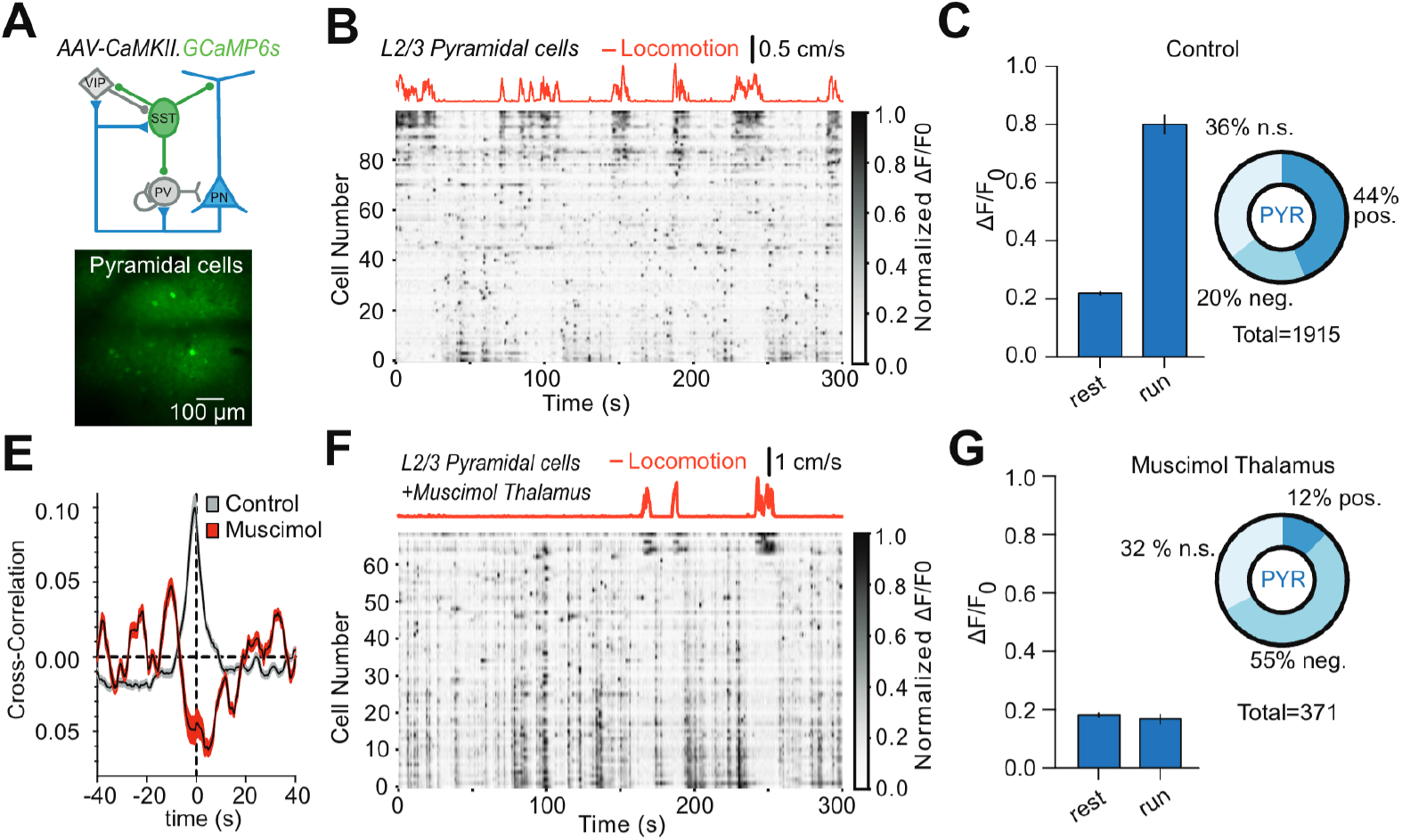
Behavioral state transitions from rest to active states increase activity of pyramidal cells in S1. A) Top: schematic illustration of layer 2/3 microcircuits in S1; bottom: in vivo 2PSLM images of S1 PNs labeled with GCaMP6s. B) Raster plot of the pyramidal cell activity in the S1 from awake mice spontaneously transitioning between resting periods and periods of locomotion. Each row represents a neuron sorted by weight on the first principal component (PC) of their activity. C) Mean ΔF/F_0_ during periods of rest and run (Control: ΔF/F_0_ rest = 0.21 ± 0.01; ΔF/F_0_ run = 0.80 ± 0.04; n = 1918 cells in 3 mice, p<0.0001, Mann-Whitney U test). E) Cross-correlation between in vivo calcium activity and running speed for Pyramidal neurons in S1 in control condition and after muscimol injections in somatosensory thalamus. F-G) Same as B and C but mice previously microinjected with muscimol in thalamus (Muscimol: ΔF/F_0_ rest = 0.18 ± 0.01; ΔF/F_0_ run = 0.17 ± 0.16; n = 371 cells in 2 mice, p<0.0001, Mann-Whitney U test).

Our observations revealed a significant rise in spiking activity among L2/3 PNs in S1 during the transition from rest to running (**Figure 7B, C**). At the population level, we noted that 44% of the cells exhibited a statistically significant increase in activity, while only 20% displayed a decrease (**Figure 7A-C**). In contrast, after muscimol injection in the thalamus this average increase in activity was completely absent (**Figure 7E-G**). In fact, positive correlation between locomotion and GCaMP6s fluorescence traces was lost upon thalamus inactivation (control: Pearson: 0.11 ± 0.01: n = 1918; run = -0.07 ± 0.01; n = 371; p <0.0001, Mann-Whitney U; **Figure 7F, G**). These results are in agreement with what has recently been reported using electrophysiological approaches in the V1^23^. Furthermore, the increase in average ΔF/F_0_ values during running periods in pyramidal cells was also not present in cells where somatosensory thalamus was inactivated by muscimol injection (control: ΔF/ F_**0**_ rest = 0.27 ± 0.01; ΔF/F_0_ run = 0.22 ± 0.02; n = 371; p <0.0001, Mann-Whitney U; **Figure 7F, G**). Hence, the convergence of locomotion-induced modulation and susceptibility to thalamic inactivation in L2/3 PNs, similar to what we observed for SST-INs, further supports the hypothesis that SST-INs in S1 inherit the increased activity during active behavioral states from L2/3 PNs.

### Differential recruitment of feedforward and disinhibitory circuits can explain the diverse modulations of SST-INs activity across conditions

Finally, we attempted to reconcile the different observations of our study into a common theoretical framework. Our experimental results highlight the critical role of the thalamus-driven cortical excitation in controlling SST-INs activity (**Figure 6, 7**) while also showing that the classical disinhibitory circuit acts as a net inhibitory input on SST-INs (**Figure 4**). We thus reasoned that the competition between those two circuits could explain the diverse modulations of SST-INs across behavioral states. We designed a minimal model implementing this idea by incorporating a thalamic afference and a disinhibitory loop (see **Figure 8A** and Methods) into a recurrent spiking network model of cortical dynamics^39,40^. The biophysical parameters and connectivity matrix are presented in Table S1 and S2 respectively. This spiking network model can be decomposed into a cortical recurrent-amplification network (performing a non-linear amplification of thalamic inputs, **Figure 8B**) and a disinhibitory circuit (performing SST-INs inhibition from VIP-INs inputs, **Figure 8B**). The network receives stationary background excitation (cell-type specific, see Table S2) as well as two time-varying inputs shaping its dynamical variations: a neuromodulatory and a sensory signal. VIP-INs are driven by the external neuromodulatory input while thalamic excitatory cells are driven by both sensory and neuromodulatory inputs (as arousal/movement-related signals also strongly drive the sensory thalamus, see^23^). Importantly, SST-INs receive inhibition from VIP-INs as well as excitation from local cortical excitatory neurons. The key ingredient of this network setting is that the VIP-mediated inhibition of SST-INs dominates at lower excitatory input levels and as activity of the excitatory input increases the strong recurrent amplification takes off and outcompetes the inhibitory action of VIP- in SST-INs (**Figure 8B**).

**Figure 8:**
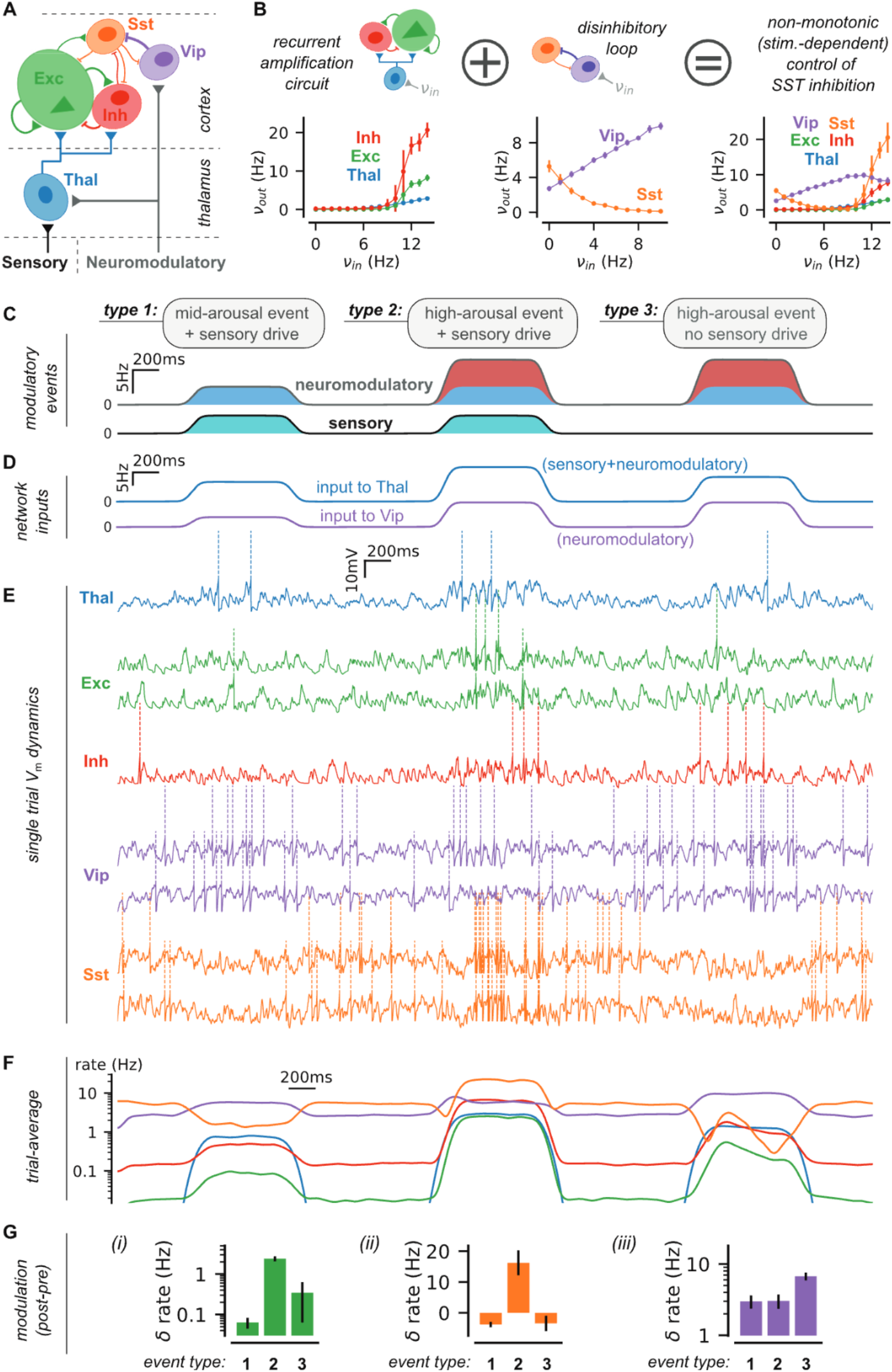
A spiking network model combining sensory and neuromodulatory inputs can explain the differential recruitment of SST-INs across conditions. **(A)** Schematic of the thalamo-cortical spiking network model (see Table S2 for connectivity parameters). The sensory input targets only the thalamus. The neuromodulatory input targets both the Vip cells and the thalamic cells. **(B)** The full network model (right) can be decomposed into a recurrent amplification circuit (left) and a disinhibitory loop (middle). We show the stationary firing levels in response to Poisson process simulations controlled by the input rate v_in_ (left, input to the thalamus only; middle, input to the Vip only; right input to both thalamic and Vip cells). Note the non-monotonic (U-shaped) response curve of Sst cells upon increasing input levels that result from the association of the two circuits (left, middle). **(C)** Modeling behavioral events through variations in neuromodulatory and sensory drives (see Methods). **(D)** Translating neuromodulatory and sensory drives into network inputs. Vip cells receive the neuromodulatory signal only. Thalamic cells receive the sum of the neuromodulatory and sensory signals. **(E)** Single trial membrane potential (*V*_*m*_) dynamics resulting from the time-varying network input of D. Note the alternations between hyperpolarizations (first and last events) and depolarizations (middle event) in Sst cells. In contrast, Vip and Exc cells are systematically depolarized. **(F)** Trial average (N=20 repeats) of the time-varying firing rate in all populations. **(G)** Firing rate modulations of the Exc *(i)*, Sst *(ii)* and Vip *(iii)* populations. Shown as mean +/- s.d. over N=20 repeats. Note that the Sst population can display either inactivation or activation following a modulatory event while the Exc and Vip display a systematic activation.

Next, we modeled the different behavioral conditions observed in our study by varying the activity of the different levels of sensory and neuromodulatory inputs feeding the network (see **Figure 8C**). We considered a first type of event (type 1) characterized by a moderate increase of arousal level (*i*.*e*. neuromodulatory input) and the presence of a sensory drive (Fig S6C, left), this is analogous to the whisking-only condition in S1. The second type of event (type 2) is characterized by a large increase of arousal and the presence of sensory inputs (Fig S6C, middle), i.e. analogous to the locomotion+whisking condition in S1. The third event (type 3) is a high arousal input in absence of sensory drive, *i*.*e*. analogous t o t h e locomotion+whisking condition with whisker-trimmed in S1. We converted those events into inputs to VIP-INs and thalamic cells (**Figure 8D**), we ran the simulations of the resulting network dynamics (**Figure 8E,F**) and we analyzed the modulation of activity elicited by the diverse event types (**Figure 8G**).

We observed that type 1 events (mid-arousal with sensory drive) led to a net inhibition of SST-INs while type 2 events (high-arousal with sensory drive) led to a net excitation of SST-Ins (**Figure 8G**). This phenomenon is explained by the non-monotonic relationship between global afferent excitation and SST-INs activity in the network (**Figure 8B**). At moderate input, SST-INs are primarily inhibited because VIP-mediated inhibition is the dominant force given the low level of cortical PN activity. At higher input levels, the thalamic drive is amplified by the cortical network and the resulting excitatory input onto SSTs (via PNs to SST-INs connections) largely exceeds the VIP-mediated inhibition. Because thalamic activity is also sensitive to neuromodulation and whisking-only events are associated with lower arousal compared to whisking+locomotion events (^41^; and therefore a reduced neuromodulatory drive) this phenomenon may provide a plausible rationale for the distinct modulations observed in SST-INs in S1 under these two different behavioral states (**Figure 2**). Similarly, in absence of sensory drive (type 3), a high arousal event is not able to bring the network at the level where recurrent amplification drives SST-INs activity (**Figure 8G**, type 3), thus potentially explaining the reduced SST-INs modulation after whisker trimming (**Figure S3**). Interestingly, those event types can also be mapped to modulatory events in V1. Indeed, type 2 would correspond to a running event (a high arousal event) in the “light” condition (i.e. in presence of sensory drive) and type 3 would correspond to a running event in the “dark” condition (i.e. no sensory drive). Here again, only in the presence of both the sensory drive and the high neuromodulatory drive, the cortical activity is able to drive SST-INs toward a strong positive modulation (**Figure 8G**, type 1 vs type 3), hence potentially explaining the observed differences in SST-INs modulation across experimental conditions (^19^ and **Figure 5**).

## Discussion

Using in vivo two-photon calcium imaging in awake mice we observed that behavioral state transitions do not equally affect the activity of L2/3 SST-INs across primary sensory regions. Specifically, we noted that in dark conditions, locomotion led to an increased activity of SST-INs in S1 but had no effect in V1. Interestingly, when visual stimulation was present, this difference was lost and transitions from rest to running now positively modulated SST-INs in both primary sensory regions. Variable recruitment of VIP-INs does not seem to be at the origin of such differential behavior-state dependent modulation of SST-INs between the two sensory regions. In fact, using a chemogenetic approach, we revealed that VIP-INs activity has a net inhibitory impact on SST-INs. In contrast, we observed that inactivation of somatosensory thalamus strongly reduced the positive modulation of SST-INs in S1 by spontaneous behaviors. Altogether our results provide a detailed description of the neuronal mechanisms at the origin of the differential modulation of SST-INs by behavior states in S1.

Our findings indicate that in S1, SST-INs are strongly recruited by locomotion even in the absence of light. This result is unexpected given the inhibitory connection between VIP-INs and SST-INs, as well as the established role of VIP-INs in inhibiting SST-INs during whisking in stationary animals^5,6,20,27,28^. Interestingly, when we selected from our recording sessions periods of whisking with no associated locomotion, we reproduced previous findings observed in non-locomoting mice. Namely, whisking resulted in an increase of VIP-INs activity and a decrease in recruitment of SST-INs^6,7^. Hence, our observations indicate that whisking during rest or during locomotion are associated with distinct cortical states, with the latter resulting in significantly stronger recruitment of SST-INs in S1.

Recent theoretical analysis has proposed a hypothesis suggesting that, somewhat counterintuitively, VIP-INs may contribute to the elevation of SST-INs activity during locomotion^30^. This phenomenon, termed “response reversal”, occurs when the increase in local excitation to SST-INs (generated from disinhibition of PNs) overcomes the VIP-mediated inhibition of SST-INs. However, we found that inhibiting VIP-INs does not abolish the positive modulation of SST-INs by locomotion; instead, it has the opposite effect, increasing the sensitivity of SST-INs to transitions from rest to running. This suggests that positive recruitment of SST-INs involves mechanisms independent of VIP-INs. Furthermore, we noted that despite relatively similar recruitment of VIP-INs across sensory regions, SST-INs in S1 and V1 displayed different response profiles to locomotion particularly in dark conditions. This result further argues that VIP-INs are likely not the cause of SST-INs increase during locomotion in the neocortex. In contrast, our experimental results illustrate that in the presence of visual stimulation modulation of SST-INs by locomotion is altered in V1 but not in S1 suggesting that sensory region-specific mechanisms are involved in such modulation.

Based on our current findings, we propose a new working model in which the positive modulation of SST-INs activity across behavioral states requires thalamic activity. Our hypothesis is strongly supported by our data, which demonstrates that inactivation of the thalamus significantly diminishes the modulation of SST-INs in S1 by locomotion. We propose that the involvement of thalamic activity partially explains the region-specific effects of spontaneous behaviors on SST-INs due to different levels of ongoing sensory drive. Activity of thalamocortical VPM neurons, that relay somatosensory information to S1, increases with whisking^42–44^. While comparatively less explored, recent findings suggest a notable upsurge in VPM activity also during locomotion^45^. It is unclear if this representation of self-motion in thalamic neurons is due to cortical inputs^43^, neuromodulation^32^ or proprioception^31^. Nonetheless, we anticipate that the combined effects of whisking and locomotion lead to an overall increase in thalamic activity, which is subsequently transmitted to the neocortex. Interestingly, our findings align with recent research indicating that depolarization of L2/3 principal neurons (PNs) in V1 during locomotion is dependent on thalamic function^23^. In fact, we obtained similar results in our study (**Figure 6**). Under normal conditions, locomotion leads to an overall increase in activity (positive LMI) in L2/3 PNs (**Figure 7**), and this increase is entirely abolished when thalamic activity is disrupted. As SST-INs are highly contacted by L2/3 PNs we suggest that the increase in SST-INs is inherited from L2/3 PNs. It is worth noting that in V1 the differential response of SST-INs to locomotion is also observed in L2/3 PN whose LMI is significantly larger in the presence of visual stimulation^19^, compatible with the idea that SST-INs display behavior dependent modulation in part transmitted from local L2/3 PNs.

Nonetheless, it is important to point out that an increase in thalamic spiking does not necessarily result in heightened activity in SST-INs. Our theoretical model suggests that this phenomenon is dependent on the combined action of neuromodulatory (linked to arousal level) and thalamocortical signals. Under conditions of low neuromodulatory or low thalamocortical activity (e.g. whisking only or dark) VIP-mediated inhibition has a dominant impact in controlling SST-INs function^6,7,21^. Instead, under conditions of enhanced sensory input, potentially intensified by higher levels of arousal, the excitatory drive transmitted by the thalamus can override VIP-mediated inhibition in SST-INs. Complementing these two sets of observation we now report that whisking and locomotion actually result in increased activity L2/3 SST-INs, suggesting that the transfer of activity across the different pathways in the somatosensory system might depend on the behavioral state of the animal. Interestingly, across the three distinct experimental conditions, the recruitment of L2/3 SST-INs appeared to closely follow the activity of local principal neurons. This suggests that the positive modulation of L2/3 SST-INs by active behavioral states is probably inherited from local PNs.

Mechanistically it is now clear that both thalamus as well as neuromodulatory inputs like cholinergic projections originating from the basal forebrain participate in the modulation of cortical networks by behavioral state. Less obvious is the computational role of the two different pathways. The cholinergic pathway, via VIP-INs activation, has been proposed to underlie the gain control of L2/3 PNs to sensory evoked activity across behavioral states^4^. Thalamic activity is required for the important depolarizations of L2/3 neurons^23^ during locomotion but their precise computational role is still ill defined. In any case by integrating feedforward thalamic activity transmitted indirectly via L2/3 PNs as well as neuromodulatory information both via VIP-mediated inhibition and via direct cholinergic innervation, SST-INs occupy a key position in cortical networks to adjust activity of the two pathways and adjust neuronal activity in function of behavior state.

### Limitations of the study

While our study points to the critical role of the sensory thalamus in indirectly controlling the modulation of SST-INs, we were unable to narrow down the specific nuclei of the somatosensory thalamus involved in such modulation because of the muscimol spread. The question of whether distinct nuclei, such as POM and VPM, make unique contributions to the modulation of SST-INs during locomotion remains unanswered.

Our study proposes a new working model where feedforward signals transmitted to the cortex by the thalamus play a critical role in determining the sensitivity of SST-INs during behavioral state transitions. However, this conclusion was primarily drawn by silencing the thalamus, as we did not directly record thalamic activity to correlate it with SST-INs modulation. Future work is therefore necessary to refine the validation of our working model.

Finally, our findings indicate that SST-INs are equipped with cholinergic receptors. While no differences were observed in the functional expression of such receptors between SST-INs in V1 and S1, we did not directly test their potential contribution to the in vivo activity of SST-INs. Therefore, we cannot exclude a possibe role of the cholinergic system in the in vivo positive modulation of SST-INs during active behavioral states in S1.

## Methods

### ^**^ Mice

Animals were housed in the Paris Brain Institute accredited by the French Ministry of Agriculture. All experiments were performed in compliance with French and European regulations on care and protection of laboratory animals (EU Directive 2010/63, French Law 2013-118, February 6th, 2013), and were approved by local ethics committees and by the French Ministry of Research and Innovation (authorization numbers #11199). All animal work was performed at the ICM PHENO-ICMice facility. Both male and female mice aged 4 weeks or older were used with the following genotypes: SST-IRES-Cre (SSTtm2.1(cre)Zjh/J; JAX 013044 ) ; SST-IRES-Cre X Ai9 (Gt(ROSA)26Sortm9(CAG-tdTomato); JAX 007909); VIP-IRES-Cre (Viptm1(cre)Zjh/AreckJ; JAX 031628) X Ai9 (Gt(ROSA)26Sortm9(CAG-tdTomato); JAX 007909); VIP-cre::SST-FLP (Ssttm3.1(flpo)Zjh/J; JAX 028579). Animals were maintained on a 12-hour light/dark cycles with food and water provided ad libitum. Animals were housed in groups of 2-6 animals per cage, as often as possible, to favor social interactions, a condition that prevents the emergence of depressive and anxious-like behavior (Berry et al., 2012; Ieraci et al., 2016; Kappel et al., 2017).

### ^**^ Slice preparation

Acute parasagittal slices (320 m) were prepared from adult mice, starting from postnatal days 25–70. Mice were deeply anesthetized with a mix of ketamine/xylazine (mix of i.p. ketamine 16 mg/kg and xylazine 13 mg/kg) and perfused transcardially with ice-cold cutting solution containing the following (in mM): 220 Sucrose, 11 Glucose, 2.5 KCl,1.25 NaH2PO4, 25 NaHCO3, 7 MgSO4, 0.5 CaCl2. After the perfusion the brain was quickly removed and slices prepared using a vibratome (Leica VT1200S). Slices containing S1 barrel field or V1 were transferred to ACSF solution at 34°C containing (in mM): 125 NaCl, 2.5 KCl, 2 CaCl2, 1 MgCl2, 1.25 NaH2PO4, 25 NaHCO3, 15 Glucose for 15–20 min. After the period of recovery, slices were kept at room temperature for a period of 5-6 h.

### ^**^ Electrophysiology

Whole-cell patch-clamp recordings were performed close to physiological temperature (32–34 °C) using a Multiclamp 700B amplifier (Molecular Devices) and fire-polished thick-walled glass patch electrodes (1.85 mm OD, 0.84 mm ID, World Precision Instruments); 3.5–5 MOhm tip resistance. For voltage-clamp recordings, cells were whole-cell patched using following intracellular solution (in mM): 90 Cs-MeSO3, 10 EGTA, 40 HEPES, 4 MgCl2, 5 QX-314, 2.5 CaCl2, 10 Na2Phosphocreatine, 0.3 MgGTP, 4 Na2ATP (300 m Osm p H adjusted to 7.3 using Cs OH). Extracellular synaptic stimulation was achieved by applying voltage pulses (20 us, 5–50 V; Digitimer Ltd, UK) via a second patch pipette filled with ACSF and placed 20–40 mm from soma. For current clamp experiments patch pipettes were filled with the following intracellular solution (in mM): 135 K-gluconate, 5 KCl, 10 HEPES, 0.01 EGTA, 10 Na2phosphocreatine, 4 MgATP, 0.3 NaGTP (295 mOsm, pH adjusted to 7.3 using KOH). The membrane potential (Vm) was held at -60 mV, if necessary, using small current injection (typically in a range between -50 pA and 200 pA). Recordings were not corrected for liquid junction potential. Series resistance was compensated online by balancing the bridge and compensating pipette capacitance. APs were initiated by brief current injection ranging from 1000–2000 pA and 1–5 ms duration. For all experiments data were discarded if series resistance, measured with a 10 mV pulse in voltage clamp configuration, was >20 MOhm or changed by more than 20% across the course of an experiment. For current clamp experiment cells were excluded if input resistance varied by more than 25% of the initial value. Recordings of puff evoked cholinergic responses in SST- and VIP-INs were performed at the resting membrane potential in the presence of 10 M CNQX and 10 M bicuculine.

All recordings were low-pass filtered at 10 kHz and digitized at 100 kHz using an analog-to-digital converter ( model NI USB 6259, National Instruments, Austin, TX, USA) and acquired with Nclamp software (Rothman and Silver, 2018) running in Igor PRO (Wavemetrics, Lake Oswego, OR, USA). The intracellular solution used for the experiments in which a presynaptic cell was depolarized by a train of action potentials to induce neurotransmitter release on a post-synaptic cell/ element was (in mM): 130 K-MeSO3, 4 MgCl2,10 HEPES, 0.01 EGTA, 4 Na2ATP, 0.3 NaGTP (300 mOsm, pH 7.3 adjusted with NaOH). Paired recordings connections were probed using a train of 5/9 action potentials at frequency of 20 Hz.

### ^**^ Surgery

Mice were initially anesthetized with Isoflurane. Induction of anaesthesia was performed at 4% isoflurane, Iso-Vet, and an air flow of 250ml/min. Mice were then placed in a stereotactic device (Kopf) kept at 37 ° C using a regulated heating blanket and a thermal probe and maintained under anaesthesia using 1-2% Isoflurane and an air flow of 250 ml/min until the end of the surgery. For calcium imaging in VIP- and SST-INs, cranial window implantation and AAV viral stereotaxic injection was performed in the same surgical procedure. In order to prevent brain swelling and post-operative pain, anti-inflammatory (Dexamethasone, 2 *μ*g/g Dexadreson, MSD) and analgesic drugs (Buprenorphine, 0.1 mg/kg Buprecare) were injected subcutaneously. After shaving the scalp, the skin was cleaned by wiping multiple times with an antiseptic solution and 70% alcohol. After injecting a local anaesthetic under the scalp (lidocaine, 10mg/kg - Xylovet or Laocaine) a section of the skin (a circular section with 1.5 cm diameter) was removed using surgical scissors. The periosteum was then carefully removed and the skull was scrapped using a dental drill, in order to remove all residues. The surface of the skull was subsequently thoroughly cleaned and dried using cottons swabs and a physiological saline solution. Using the stereotactic device, the center of the cranial window was marked and a circular piece of bone was removed using a 3 mm biopsy punch (LCH-PUK-30 Kai Medical). This step of the surgical procedure was considered critical since damage to the dura would result in rapidly opacifying cranial windows. Once the brain was exposed, the AAV viral vectors carrying the genetically encoded calcium indicators (GCaMP6 f and s) were stereotactically injected. For calcium imaging of VIP-INs, SST-INs and pyramidal neurons, AAV9 Syn-Flex-GCaMP6f-WPRE-SV40 (2.1x1013 AddGene-100833), AAV9 Syn-Flex-GCaMP6s-WPRE-SV40 (2.5x1013 AddGene-100845) or AAV9.Syn.GCaMP6s.WPRE.SV40 (2.7x1013 AddGene-100843) was used respectively. Using a Nanojet III (Drummond Scientific), 200 nL of virus were injected in the barrel cortex (from Bregma, 1.1 RC; 3.3 ML) or visual cortex (from bregma: 3.0 RC; 2.4 ML) at a rate of 1 nL/s followed by 5 minutes waiting period to prevent backflow. A round of 3mm glass coverslip (CS-3R Warner Instruments) was then placed over the craniotomy and glued in place using dental cement (Superbond). A stainless-steel head post (Luigs & Neumann) was then attached to the skull also using dental cement. For 3 days following the cranial windows surgery, mice were injected once a day with a mix of anti-inflammatory (Dexamethasone, 2 *μ*g/g Dexadreson, MSD) and analgesic drugs (Buprenorphine, 0.1 mg/kg Buprecare). Two weeks after surgery, mice were first habituated to the experimenter by handling and the following days, mice were head-fixed in the acquisition setup for increasing periods of time (5 – 45 minutes). The habituation of animals to the experimental setup lasted at least 10 days with a minimum of 5 sessions. At the end of the habituation period animals spontaneously transitioned between rest and running periods in the circular treadmill. Animals were maintained in a normal light-dark cycle and imaged between Zeitgeber time 0 and 12. Imaging sessions lasted between 30 to 45 min and animals were imaged between 3-5 days.

### ^**^ Thalamus inactivation

In order to inactivate the somatosensory-associated thalamus, following the acquisition of several control imaging sessions, mice were placed in a stereotactic device, and put under light isoflurane anesthesia. A hole was drilled over the ipsilateral thalamus and using a cannula, a bolus of fluorescent Muscimol (0.5 mM, Muscimol, BODIPY&trade; TMR-X Conjugate Muscimol, BODIPY™ TMR-X Conjugate, ThermoFischer Scientific) was slowly injected into the thalamus (Coordinates: -1.82 RC, 1.5 ML, 3.5 depth). After injection, the craniotomy was covered with silicone (Kwik-Cast, WPI) and the mouse was allowed to recover for an additional 30 minutes. After recovery, two-photon imaging was performed for a maximum of one hour post muscimol injection. The effective inactivation of the thalamus was confirmed functionally by recording the response of pyramidal cells as well as local cortical interneurons to whisker stimulation as well as by performing histological analysis just after the imaging sessions. For that, following the conclusion of the experiment, mice were anesthetized with isoflurane and the brain quickly removed. Coronal brain sections with a thickness of 300 *μ*m were generated to verify the injection site and distribution of muscimol using fluorescence imaging.

### ^**^ Intrinsic Signal Imaging

Intrinsic signal imaging (ISI) was used to obtain spatial maps of visual areas in the visual cortex and thus to confirm that the injection site was located in V1 (Fig. S5). The procedure was reproduced from^46^. Briefly, mice were injected with Chlorprothixene (Santa Cruz Biotechnology) and lightly anesthetized with 0.75–1.5% isoflurane. Imaging sessions began with a vasculature image acquired under green illumination (530-nm LEDs; Thor Labs). Next, the imaging plane was defocused to 500 *μ*m below the vasculature plane. The haemodynamic response to a visual stimulus was imaged under red light (625-nm LED; Thor Labs) with a sCMOS camera (Thorlabs, Kiralux CC505MU). The stimulus consisted of an alternating checkerboard pattern (20° wide bar, 25° square size) moving across a grey background. On each trial, the stimulus bar was swept across the four cardinal axes 10 times in each direction at a rate of 0.1 Hz. Up to 5 trials per direction were performed on each mouse. Altitude and azimuth phase maps were calculated from the discrete Fourier transform of each pixel. A ‘sign map’ was produced from the phase maps by taking the sine of the angle between the altitude and azimuth map gradients. V1 was then easily identified as the constant sign area with a high intrinsic signal response (Fig. S5).

### ^**^ Two-Photon Imaging

In vivo two-photon calcium imaging was performed with an Ultima IV two-photon laser-scanning microscope system (Bruker), using a 20X, 1.0 N.A. water immersion objective (Olympus) with the femtosecond laser (MaiTai DeepSee, Spectra Physics) tuned to 920 nm for imaging of cells expressing GCaMP6s/f and to 1020 nm for imaging of cells expressing mCherry. Fluorescence light was separated from the excitation path through a long pass dichroic (660dcxr; Chroma, USA), split into green and red channels with a second long pass dichroic (575dcxr; Chroma, USA), and cleaned up with band pass filters (hq525/70 and hq607/45; Chroma, USA). Fluorescence was detected using both proximal epifluorescence photomultiplier tubes (gallium arsenide phosphide, H7422PA-40 SEL, Hamamatsu). Time-series movies of neuronal populations expressing GCaMP6f were acquired at the frame rate of 30Hz (512 × 512 pixels field of view; 1.13 *μ*m/pixel). The duration of each focal plane movie was 300s (9000 frames) to track spontaneous neuronal activity. During the recording periods animals were free to run on a circular treadmill covered with a soft foam. This foam softened the movement of the animals and provided better traction (this was important since mice adapted more rapidly to the experimental setup and movements were more fluid during the recordings). Locomotion was monitored using a rotary encoder (H5-1000-NE-S, US Digital) and digitized with a NIDAQ analog-to-digital converter (model NI USB 6393, National Instruments, Austin, TX, USA) at 2000 Hz. Whisker movement was recorded using a FLIR camera (BFS-U3-17S7M-C) at a frame rate of 30Hz equipped with a cleanup filter (FB850-40, Thor labs) to block laser light from two-photon imaging and controlled via the Spinnaker SDK software (FLIR Systems Inc, Orlando, USA). The camera was focused on the whisker pad of the mouse using a 10X (13 - 130mm FL) C-Mount, Close Focus Zoom lens and each frame acquisition was controlled via Neuromatic running in IGORPRO (Wavemetrics). The mouse snout was illuminated via an 850 nm LED (M850L3, Thor labs). In a subset of mice location of imaging was evaluated using histological analysis. For that, mice were transcardially perfused with PBS 1X, followed by 4% paraformaldehyde in PBS (PFA). After extraction, brains were post-fixed overnight in PFA 4% at 4°C and preserved in PBS 1X at 4°C. To evaluate location of GCaMP6s expression brains were first dehydrated through incubation in 30% sucrose in PBS 1X overnight at 4°C, followed by freezing in methyl-butane and kept at -80°C. Coronal slicing was carried out using a freezing microtome, at a thickness of 100*μ*m. Sections were mounted on microscope slides with fluoromount aqueous mounting medium (Sigma). Images were acquired using a Zeiss ApoTome.2 microscope with 5X and 10X objectives.

### ^**^ Whisker Stimulation

For whisker stimulation experiments, a NPI PDES-02DX (NPI Electronic GmbH) puffer was used. Tubing was connected to the puffer and a short puff (1 sec) of air was aimed at the mouse whiskers. The tip of the puffer was kept out of reach of exploratory whisking range. The whisker puff was randomly triggered during imaging sessions and the time of puff was recorded via the digitization of voltage trigger using a analog-to-digital converter (model NI USB 6259, National Instruments, Austin, TX, USA)

### ^**^ CNO Experiments

For experiments involving DREADDs, mice were stereotactically injected in S1 with the AAV viral vector carrying the hM4D(Gi) receptor together with the reporter mCherry. The hM4D(Gi) receptor was selectively expressed in VIP-INs by injecting AAV9 . hSyn . DIO . hM4D ( Gi ) - mCherry (AddGene-44362), in VIP-Cre mice. Mice were used for experiments at least 3 weeks after viral injection. For experiments, mice were first imaged to monitor activity of SST-INs in control conditions and subsequently intraperitoneally injected with clozapine-N-oxide (CNO). Activity of SST-INs was re-evaluated 30 min after CNO injection. A water-soluble salt, CNO dihydrochloride (Tocris, Bio-Techne LTD, Abingdon, UK, catalog no.: 6329), was used for the preparation of CNO solution, that was dissolved in sterile saline. Several concentrations of CNO were tested (1, 5 and 10mg/kg) and no significant changes were found so data was all data was pulled together.

### ^**^ Visual stimulation

To study the effect of light on the recruitment of VIP- and SST-INs in the cortex, an isoluminant grey screen (Lilliput 669GL 7”) was placed at 45° from the center line of the mouse and at approximately 20cm from the left eye (contralateral to the craniotomy) of the mouse. Visual stimulation trials were interleaved with control trials. In order to reduce light contamination, a custom funnel was used to shield the objective from light coming from the screen.

### ^**^ Calcium Imaging Analysis

Motion correction, image segmentation and calcium signal extraction were performed using the python 3.6 package Suite2P^47^. This package analyses data in four stages: image registration, region of interest (ROI) detection, quality control, and activity and neuropil extraction. The first stage, image registration, is responsible for motion correction and works by performing a mean image of a few frames and then shifting (in the x and y axis) each frame so that it optimally fits the mean image. The second stage, ROI detection, detects the cells in the frame. This is done by detecting the pixels with the highest variability across time and then expanding the region to the surrounding pixels until this variability drops below a defined threshold. Although this method works quite well, the disadvantage of this method is that since only active cells are detected, it is difficult to determine the number of silent cells in field of view, which could be important for functional tasks. In a third stage, quality control, the program separates structures that correspond to cells from the remaining structures by assigning them a score. The results of this stage were always manually curated. In the last stage of the analysis, activity and neuropil extraction, is performed by averaging the pixel intensity in the ROI for each time point.

### ^**^ Analysis of fluorescence signals

Raw fluorescence traces were corrected for neuropil contamination by subtracting Suite2p-neuropil traces using a fixed scaling coefficient of 0.7^18^. To be able to compare activity across cells and mouse lines, fluorescence signals were normalized. To do this we used the ΔF/F_0_ method, calculated using the following formula: (F-F_0_)/F_0_, where F is the fluorescence and F_0_ is the baseline fluorescence. When studying the activity of interneurons (VIP- and SST-INs), the baseline fluorescence was calculated using the lower 5th percentile. Results were systematically evaluated using a different baseline method to estimate F0 (MinMax baseline method with a sliding window of 60s) and observations reported in the present manuscript were insensitive to the baseline method used. For the analysis of pyramidal cell activity, a sliding average window (60 s) defined as MinMax method was used.

### ^**^ Locomotion Analysis

Changes in positioning of the circular treadmill (sampled at 2 kHz) were interpolated onto a down-sampled rate of 30Hz in order to match the sampling rate of the two-photon imaging. Locomotion periods were defined as the periods where speed was higher than 0.1 cm/s. All points 0.5 seconds before and 1.5 seconds after a locomotion period were considered stationary periods. Events spaced by less than 0.5s were joined. Sessions in which the percentage of either locomotion or stationary periods were lower than 5%, were excluded. The locomotion modulation index (LMI) was defined as the difference between the mean ΔF/F0 during locomotion (F_run_) and stationary (F_rest_) periods, normalized by the sum of both values: LMI = (F_run_ – F_rest_)/( F_run_ + F_rest_)^19^. To calculate statistically significant Pearson Correlations between fluorescence and running speed, comparisons were made with 1000 shuffled (circular shuffling) locomotion traces^18^. Pearson correlation coefficient was considered statistically significant if the value of original data was above a 95% confidence interval generated from the 1000 shuffled traces.

### ^**^ Whisking Analysis

Whisker pad movement was tracked using the python framework Facemap^48^. Whisker-pad movement was measured by selecting an ROI over the whisker pad and calculating the motion energy index (MEI) across the video. MEI was defined as the sum of the absolute change in pixel intensity within the ROI between adjacent video frames. Using custom python code, whisking pad motion energy was filtered with a gaussian filter (sigma of 3), normalized with a Min Max method and resampled to 30Hz. Whisking periods were defined as periods where normalized MEI exceeded 10 %. Whisking only periods were defined as periods in which there was whisking but not locomotion. Whisking modulation index (WMI) and whisking only modulation index (WOMI) were calculated in a similar way to LMI but using the variations in GCaMP6s fluorescence during resting or whisking periods.

Event-triggered averages were computed by automatically identifying frames with event onsets, namely those related to whisking and/or locomotion. Due to the slow time course of GCaMP6s and to minimize cross-talk between events, we specifically chose locomotion events that were not preceded by another locomotion period within a 5-second interval. For whisking only analysis, whisking episodes not associated with locomotion were included in the average (5s before and 5s after). In Fig. 3I, locomotion events were further categorized in agreement with locomotion duration. Once multiple locomotion and/or whisking events were detected, the results were combined and averaged to calculate an average value for each cell. Subsequently, these individual average cell values were averaged together and presented with their respective average and SEM.

### ^**^ Spiking Network Model

We analyzed the modulation evoked by diverse neuromodulatory and sensory events in a numerical model of neural network dynamics: a randomly connected network of artificially-spiking neurons with conductance-based synapses ^39,40^. Single cells were described as single compartment leaky integrate-and-fire models and the synaptic dynamics followed an exponential time course. The dynamics of a single neurons thus followed the set of equations:

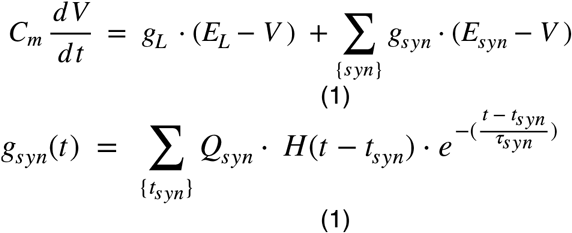

where *V*_*m*_ is the membrane potential of the neuron and *C*_*m*_ its membrane capacitance. The leak currents were set by the leak conductance *g*_*L*_ and leak reversal potential *E*_*L*_. The synaptic currents (indexed by ‘‘syn’’ and summing over its set of source synaptic population for a given neuron type) were set by the time-varying synaptic conductances *g*_*syn*_*(t)* and the synaptic reversal potential *E*_*syn*_. The synaptic conductances *g*_*syn*_*(t)* was given by the convolution of the set of incoming synaptic events {*t*_*syn*_} with an exponential waveform of time constant τ_syn_ weighted by the synaptic quantal Q_syn_. The membrane potential dynamics (Equation 1) was complemented with a ‘‘threshold and reset’’ mechanism. When the membrane potential reached the spiking threshold, it was clamped at the reset potential for the duration of the refractory period after which the *V*_*m*_ dynamics restarted from there. The cellular parameters can be found in the table S1.

The network is made of 5 populations: an excitatory thalamic population (indexed “Thal”), two cortical populations forming the core excitatory/inhibitory network behind the non-linear recurrent amplification phenomenon (4000 and 800 neurons, indexed by “Exc” and “Inh” respectively), the VIP-INs inhibitory population (100 neurons, indexed “VIP”) and the SST-INs inhibitory population (100 neurons, indexed “SST”). An external excitatory population of 200 neurons (indexed by “BgExc”) firing with Poisson statistics at a constant rate of 20 Hz provides some background excitatory drive to the network. Two external populations of 100 neurons targeting the thalamic cells and the Vip cells (indexed “ThalExc” and “VipExc” respectively) control the time-varying dynamics of the network. Those two populations are assigned a time-varying rate (see Fig. S6D) that is translated into discrete spiking events through an inhomogeneous Poisson process. The connectivity matrix between all populations can be found in Table S2.

To model behavioral events, the levels of neuromodulatory and sensory inputs were adjusted to fit within the range of inputs where the output rate of all populations would avoid too dense firing (i.e. below 25 Hz). The corresponding range of inputs was found to be 0-14 Hz (see Fig. S6B, right). This led us to set amplitudes of events as: 4Hz for sensory drive event, 4 Hz for mid-arousal neuromodulatory event, 10 Hz for high-arousal neuromodulatory event.

Numerical simulations were performed with the Brian2 simulator^49^. A time step of dt = 0.1ms was chosen. Simulations were repeated over multiple seeds (**Figure 8**) generating different realizations of the random connectivity scheme and of the external drive.

### ^**^ Statistics

All statistical analysis was performed in Prism (GraphPad Software, Inc.). The error bars in all graphs indicate standard error of the mean (S.E.M.) and statistics were performed with two-tailed tests. For statistical tests comparing the average ΔF/F0 of neurons between two behavioral states (stationary versus locomotion periods) in the same subject, we used Wilcoxon signed-rank tests. For statistical tests comparing the distribution of LMI, Pearson Correlation or behavior properties between brain regions or interneuron populations, Mann-Whitney U tests were used. When comparing two conditions across behavioral states, Kruskal-Wallis test was used.

### ^**^ Data and code availability

All data reported in this paper will be shared by the lead contact upon request.

Custom scripts used in this study have been deposited to a publicly accessible repository. The URL is listed in the key resources table.

Any additional information required to reanalyze the data reported in this paper is available from the lead contact upon request.

## ^**^ Acknowledgements

This study was supported by the Centre National de la Recherche Scientifique, Agence Nationale de la Recherche (grants ANR-EXCIGLY and ANR-DecoSensoMol1) the European Research Council (ERC-STG-678250 to N.R.), the European Union’s Horizon 2020 research and innovation programme (Marie Skłodowska-Curie Grant 892175 “InProsMod” to Y.Z.); by the Paris Brain Institute, and the Fondation pour la Recherche Médicale (FRM; fellowship ARF201909009117 to Y.Z.). We thank the Paris Brain Institute core facilities, namely iVector and ICMice PHENOPARC. We would also like to thank Dr. Alberto Bacci (ICM, Paris) and Dr. Joana Lourenco (ICM, Paris) for the valuable discussions and comments on the manuscript.

---

**Supplementary Fig. 1:**
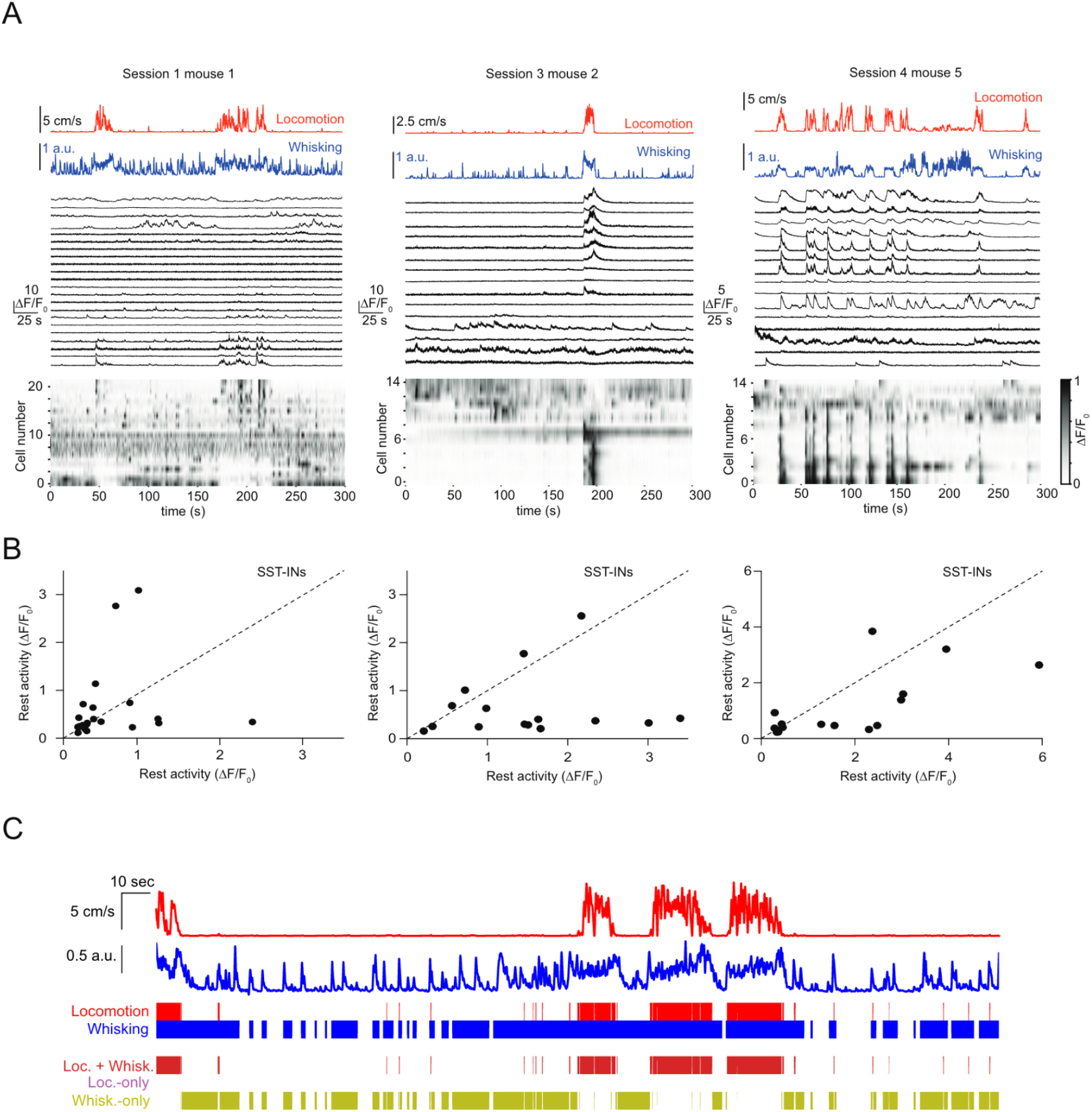
Variability of SST-INs response within individual fields of view in S1 to behavioral transitions from rest to run. A) Representative GCaMP6s fluorescence traces (black) shown as ΔF/F_0_ (up) as well as normalized ΔF/F_0_ (bottom) for SST-INs for individual fields recorded in awake mice spontaneously transitioning between rest and run periods. Each row represents a neuron sorted by weight on the first principal component (PC) of their activity. Locomotion (red) and whisking (blues) traces are shown on top. B) Scatter plots of the mean amplitude of fluorescence changes (ΔF/F_0_) of each neuron illustrated in A for locomotion periods versus stationary periods. C) Example session, in addition to the one in Figure 2A, illustrating simultaneous recording traces for whisking (blue) and running speed (red). Whisking only periods (no locomotor activity) are highlighted in light green. Note in this session the higher (compared to the one of Figure 2A) proportion of whisking only periods.

**Supplementary Fig. 2:**
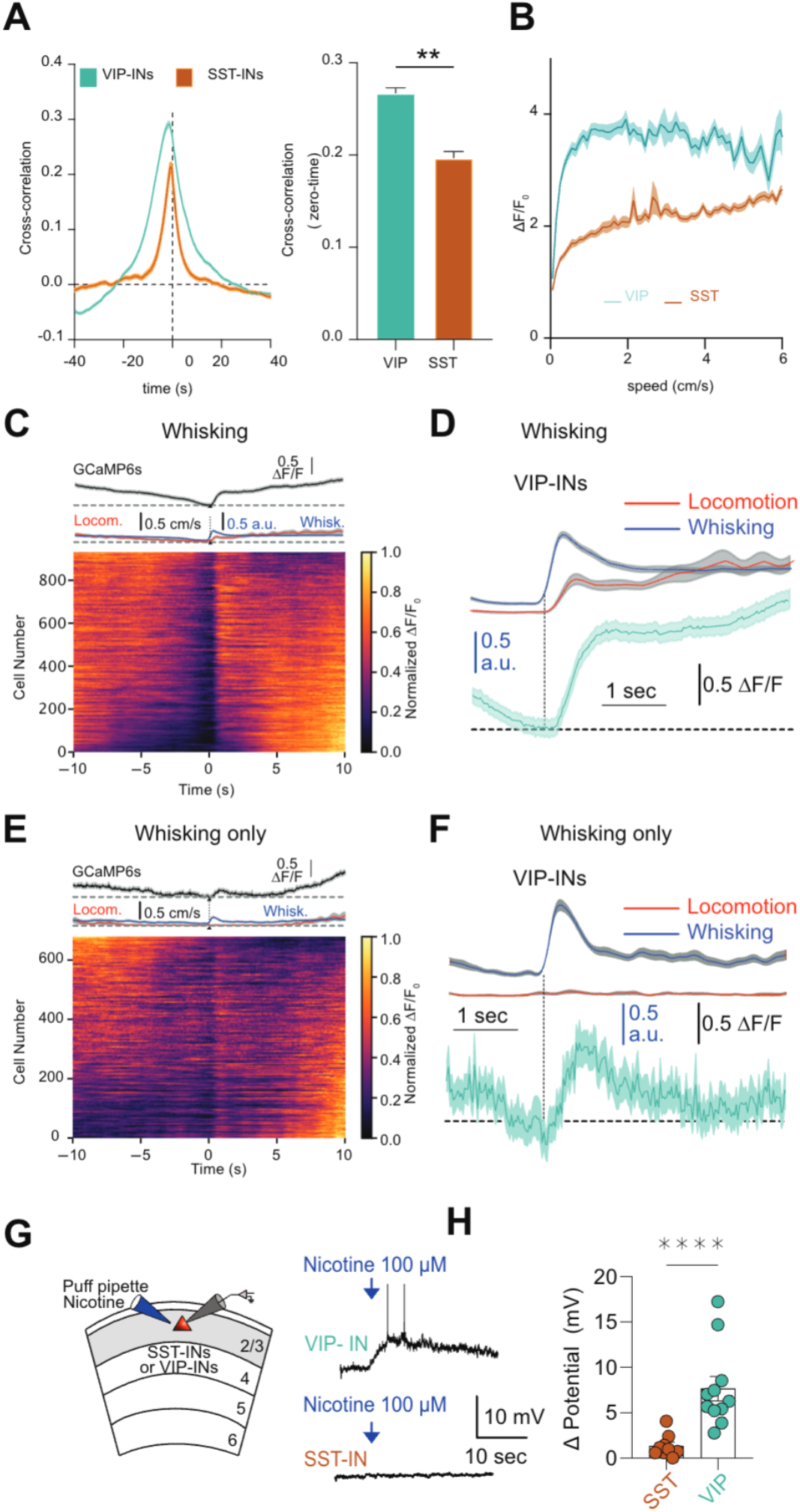
Response of SST- and VIP-INs in S1 to behavioral transitions from rest to run. A) Cross-correlation curves between running speed and calcium activity for VIP and SST-INs. Values at zero time are displayed on the right. B) Variation in ΔF/F values for SST- and VIP-INs with average running speed. C) Event-triggered average traces of VIP-INs GCaMP6s fluorescence signal in S1 aligned to whisking onset. Normalized ΔF/ F_0_ is displayed for all analyzed cells (bottom). Traces represent mean ± SEM values across all recorded cells. (D) Enlarged view of average traces shown in C (top). E, F) Same as C and D but for whisking only periods. G) Schematic illustration of whole-cell current clamp recording in L2/3 of primary somatosensory (S1) cortex while puff applying nicotine 100 nM in SST or VIP-INs (left). Example traces of puff-evoked responses in SST- and VIP-INs (right). H) Summary plot of variations in membrane potential induced by nicotine applications in both SST- and VIP-INs (SST [n = 11] vs VIP [n = 9], p < 0.0001, Mann-Whitney test).

**Supplementary Fig. 3:**
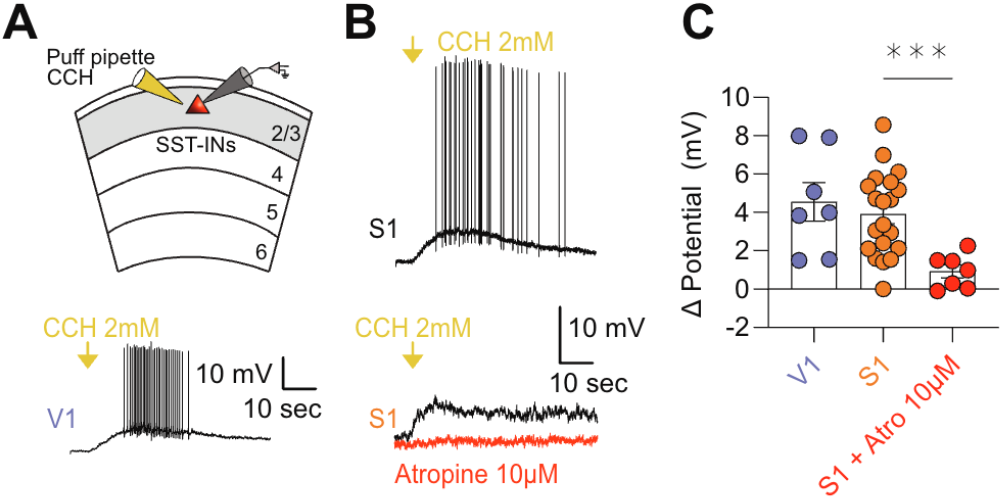
SST-INs are equipped with cholinergic receptors in both V1 and S1. A) Schematic illustration of whole-cell current clamp recording in L2/3 of primary somatosensory (S1) cortex while puff applying carbachol (2mM) in SST (up). Example traces of puff-evoked responses in SST-INs (down). B) Example traces of current clamp recordings illustrating the variation in membrane potential in SST-INs to puff application of carbachol in both V1 (P>56: top right) and S1 (P<56 middle right; P>56 bottom right). C) Summary plot of the variation of membrane potential induced by carbachol (CCH) in SST-INs in V1 and S1 (V1, SST [n = 7]; S1, SST with CCH puff alone [n = 20] versus SST with CCH puff in the presence of atropine 10*μ*M [n = 7], p = 0.007, Mann-Whitney test). Error bars are SEM.

**Supplementary Fig. 4:**
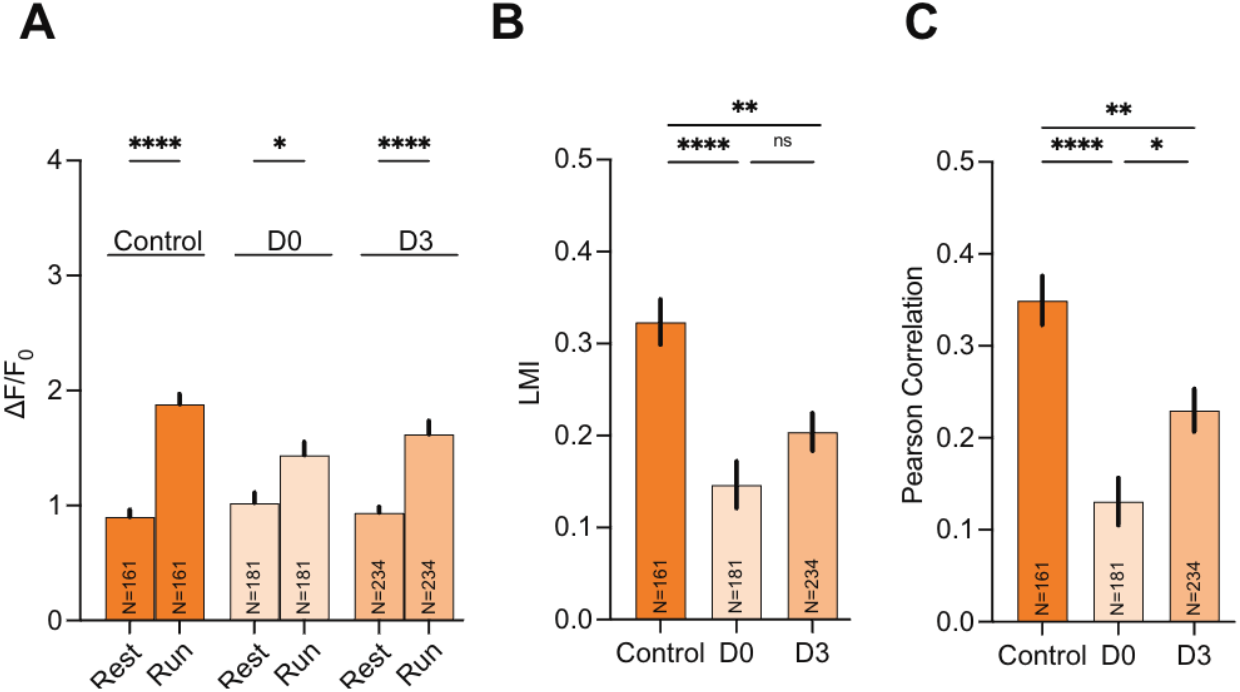
Whisker trimming transiently reduces sensitivity of S1 L2/3 SST-INs to locomotion. A) Mean ΔF/F0 of GCaMP6s in SST-INs during periods of rest and run in control conditions and during 3 days after whisker trimming. B and C) Summary plots of effect of whisker trimming in LMI and Pearson correlation value between locomotion and GCaMP6s fluorescence signal. Values represent mean ± SEM.

**Supplementary Fig. 5:**
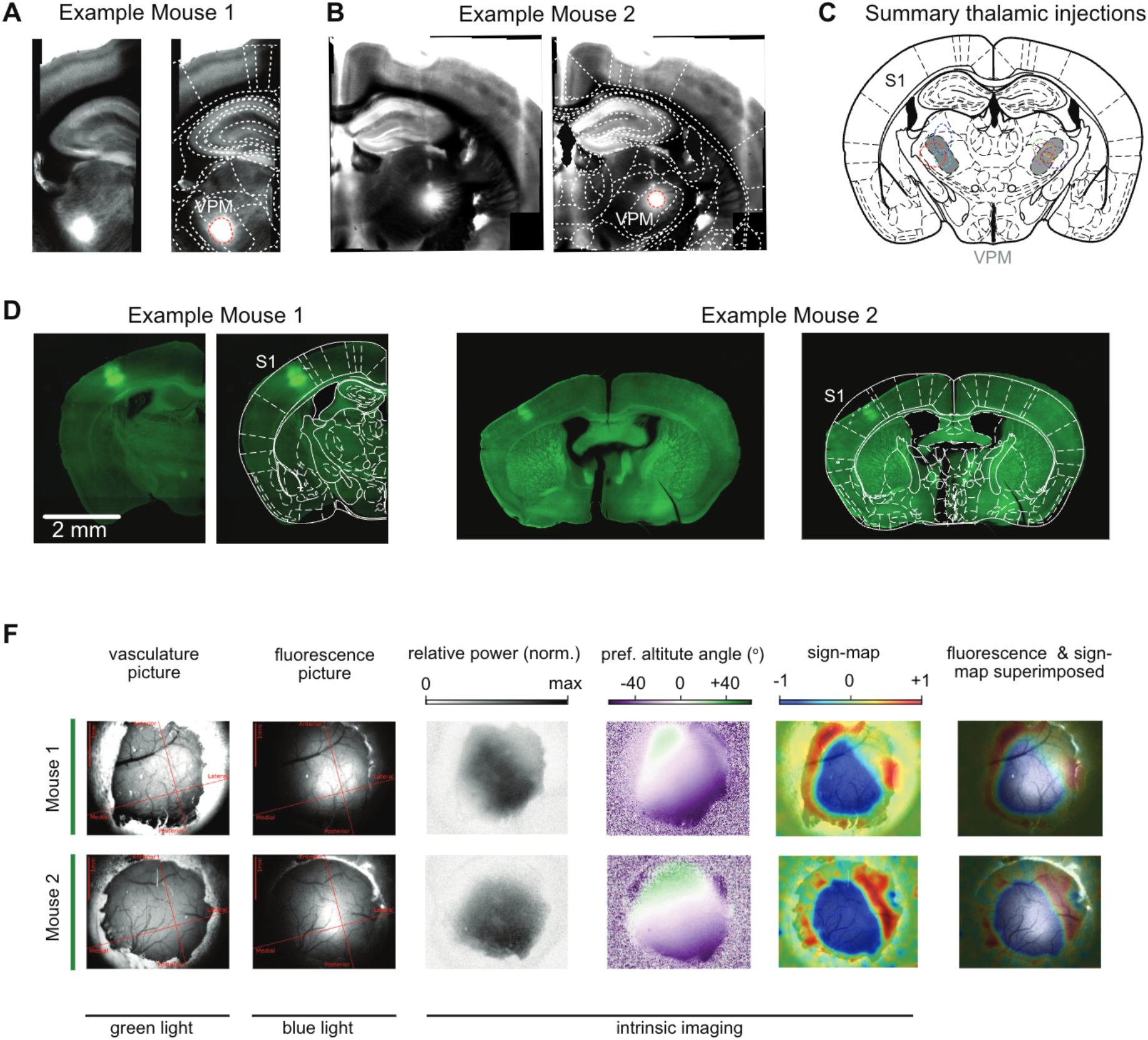
Evaluation of site of muscimol injection and validation of S1 and V1 imaging locations. A) (left) Differential interference contrast (DIC) microscopy image with merged fluorescence signal from a coronal brain slice obtained from a mouse injected with muscimol and used to obtain in data presented in Fig. 6. (right) Same as left but with superimposed image obtained from Paxinos and Franklin Atlas. B) Same as A but for a different mouse. C) Summary plot of the site of muscimol injection for the 8 mice used in the study. Dotted line represents the FWHM of the fluorescence signal. D) Example of the location of GCaMP6s fluorescence signal in two animals used to monitor in vivo activity of L2/3 PNs in S1. Note that in both cases expression correctly targets S1 Barrel cortex). F) Intrinsic optical imaging with superimposed fluorescence detection of GCaMP6s signals validates the accurate targeting of the mouse primary visual cortex.

**Table S1.** Biophysical parameters of the spiking network model.

| Name | Symbol | Value |
| --- | --- | --- |
| Membrane Capacitance | $C_m$ | 200 pF |
| Leak Conductance | $g_L$ | 10 nS |
| Leak Reversal Potential | $E_L$ | -70 mV |
| Spike Reset Potential | $V_{reset}$ | -70 mV |
| Refractory Period | $\tau_{refrac}$ | 5 ms |
| Spike Threshold (Thal, Exc) | $V_{thre}^{Exc}$ | -50 mV |
| Spike Threshold (Inh, Sst, Vip) | $V_{thre}^{Inh}$ | -53 mV |
| Synaptic Quantal (BgExc, Thal) | $Q_{exc}^1$ | 4 nS |
| Synaptic Quantal (Exc, VipExc, ThalExc) | $Q_{exc}^2$ | 1 nS |
| Synaptic Quantal (Inh, Sst, Vip) | $Q_{inh}$ | 10 nS |

**Table S2.**
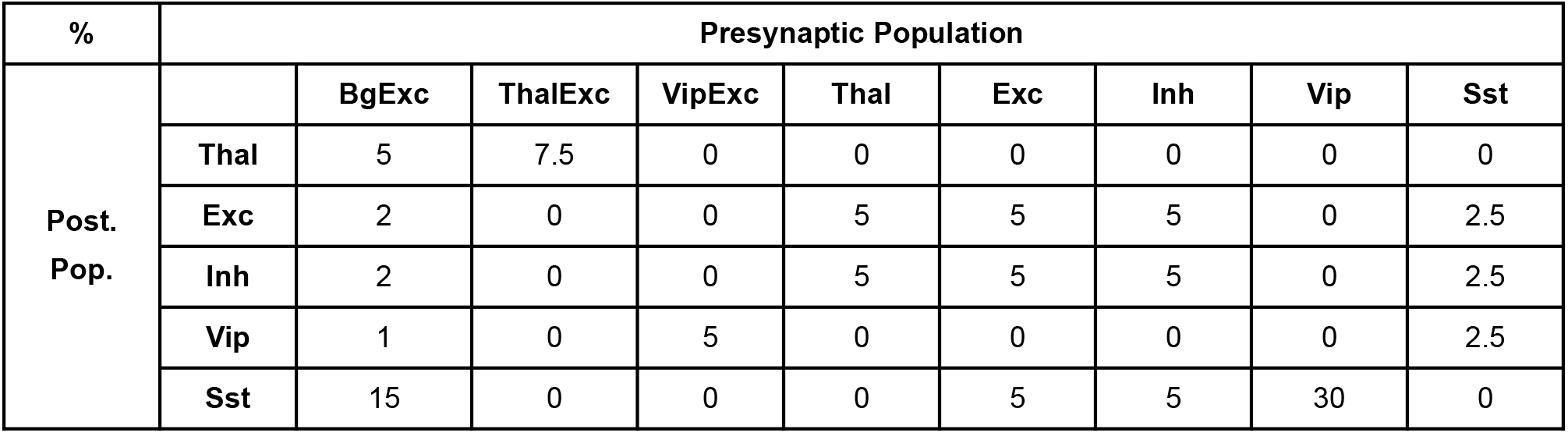
Connectivity parameters of the spiking network model. Connectivity is expressed in percent of the postsynaptic population, i.e. a population X connecting to a population Y of 1000 neurons with a connectivity of 5% will correspond to randomly picking 50 target cells in the Y populations for each cell in the population X.

## Key Resources Table

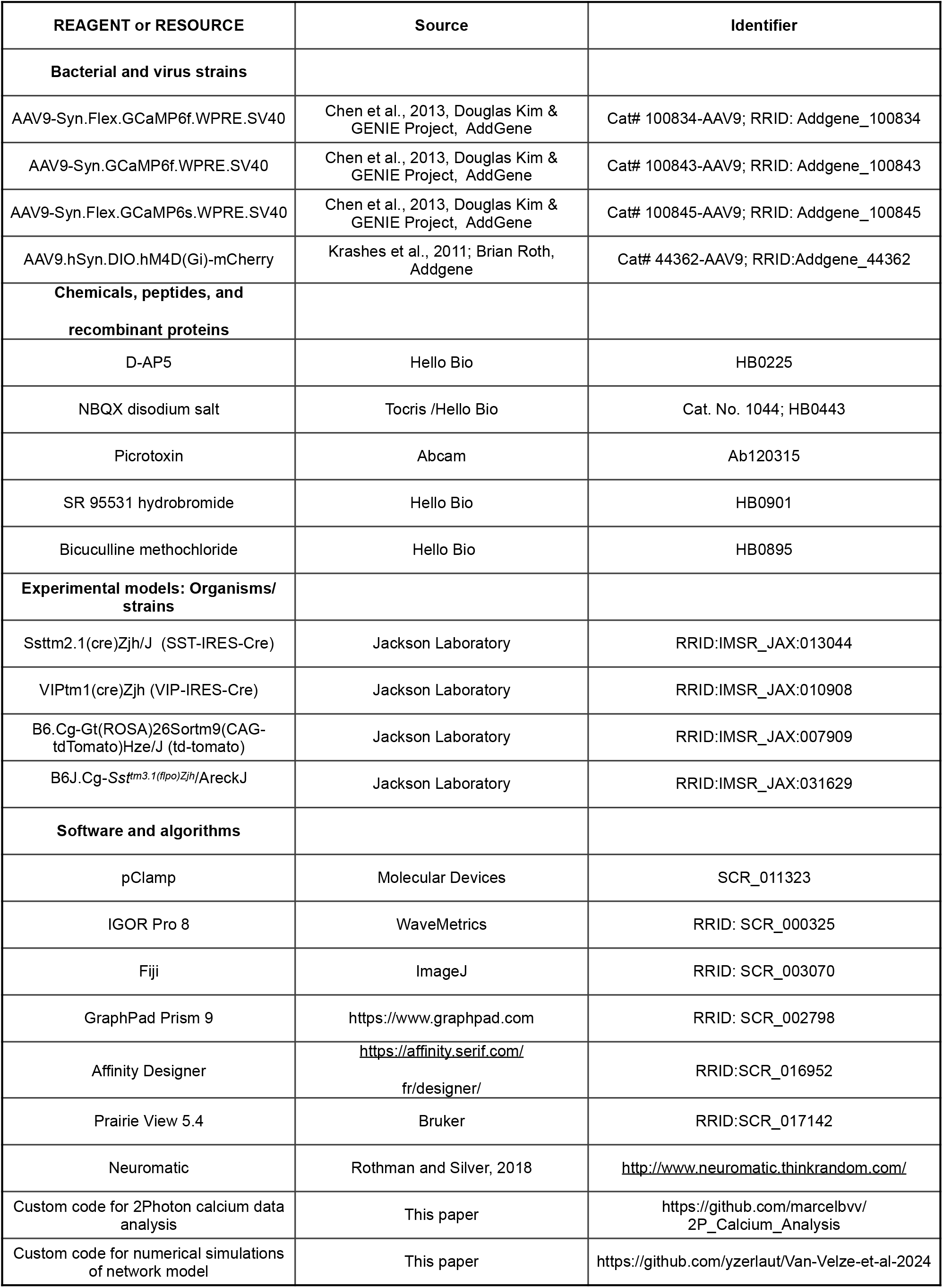

## Notes

### Competing Interest Statement

The authors have declared no competing interest.

### Summary of Updates

The title has been modified and a new result section using model simulations has been included.

